# ERBB4-mediated signaling is a mediator of resistance to BTK and PI3K inhibitors in B cell lymphoid neoplasms

**DOI:** 10.1101/2023.01.01.522017

**Authors:** Alberto J. Arribas, Sara Napoli, Luciano Cascione, Laura Barnabei, Giulio Sartori, Eleonora Cannas, Eugenio Gaudio, Chiara Tarantelli, Afua A. Mensah, Filippo Spriano, Antonella Zucchetto, Francesca M. Rossi, Andrea Rinaldi, Manuel Castro de Moura, Sandra Jovic, Roberta Bordone Pittau, Anastasios Stathis, Georg Stussi, Valter Gattei, Jennifer R. Brown, Manel Esteller, Emanuele Zucca, Davide Rossi, Francesco Bertoni

## Abstract

BTK and PI3K inhibitors are among the drugs approved for the treatment of patients with lymphoid neoplasms. Although active, their ability to lead as single agents to long-lasting complete remission is rather limited especially in the lymphoma setting. This indicates that tumor cells often develop resistance to the drugs. Here, we show that the overexpression of ERBB4 and its ligands represents a modality for B cell neoplastic cells to bypass the anti-tumor activity of BTK and PI3K inhibitors and that targeted pharmacological interventions can restore sensitivity to the small molecules.

We started from a marginal zone lymphoma (MZL) cell line, Karpas-1718, kept under prolonged exposure to the PI3Kδ inhibitor idelalisib until acquisition of resistance, or with no drug. Cells underwent transcriptome, miRNA and methylation profiling, whole exome sequencing, and pharmacological screening which led to the identification of the overexpression of ERBB4 and its ligands HBEGF and NRG2 in the resistant cells. Cellular and genetic experiments demonstrated the involvement of this axis in blocking the anti-tumor activity of various BTK and PI3K inhibitors, currently used in the clinical setting. Addition of recombinant HBEGF induced resistance to BTK and PI3K inhibitors in parental cells but also in additional lymphoma models. Combination with the ERBB inhibitor lapatinib was beneficial in resistant cells and in other lymphoma models already expressing the identified resistance factors. Multi-omics analysis underlined that an epigenetic reprogramming affected the expression of the resistance-related factors, and pretreatment with demethylating agents or EZH2 inhibitors overcame the resistance. Resistance factors were shown to be expressed in clinical samples, further extending the findings of the study.

In conclusions, we identified a novel ERBB4-driven mechanism of resistance to BTK and PI3K inhibitors and treatments that appear to overcome it.

**Key points:**

- A mechanism of secondary resistance to the PI3Kδ and BTK inhibitors in B cell neoplasms driven by secreted factors.
- Resistance can be reverted by targeting ERBB signaling.

## Introduction

B cell receptor (BCR) signaling is a crucial pathway for the proliferation and the survival of B cells, and its pharmacological inhibition with BTK and PI3K inhibitors has proven its clinical efficacy for the treatment of patients affected by lymphoid neoplasms (1–5). The PI3Kδ inhibitor idelalisib and the BTK inhibitor ibrutinib were the two agents belonging to these classes to be first approved (2,5). They have been successfully followed by newer compounds, including second generation PI3Kδ inhibitors (parsaclisib or zandelisib), small molecules that inhibit PI3Kδ but also other kinases, such as copanlisib (PI3Kα/PI3Kδ), duvelisib (PI3Kγ/PI3Kδ) or umbralisib (PI3Kδ and casein kinase-1ε), BTK inhibitors with higher specificity for the target (zanubrutinib, acalabrutinib) or with different mechanism of action (pirtobrutinib) (2,4,5). However, single agent treatments is often associated with limited success in achieving long-lasting complete responses, suggesting the need for combination therapies (6). The identification of the mechanisms underlying either primary or secondary resistance to signaling inhibitors (5,7–15) provides useful information to optimize the use of the agents. Here, starting from a marginal zone lymphoma (MZL) cell line kept under prolonged exposure to the PI3Kδ inhibitor idelalisib, we demonstrate that resistance to BTK and PI3K inhibitors can be sustained in various B cell lymphoid neoplasms by the overexpression of ERBB4 and its ligands and targeted pharmacological interventions can restore sensitivity to the small molecules.

## Materials and Methods

### Development of resistant cell lines

Karpas1718 cells (16) were cultured according to the recommended conditions, as previously described (12). All media were supplemented with fetal bovine serum, penicillin-streptomycin-neomycin (5,000 units penicillin, 5 mg streptomycin and 10 mg neomycin/mL, Sigma) and L-glutamine (1%). To develop resistance cell lines were exposed to IC90 concentration of idelalisib (Selleckchem, TX, USA) for several months until they acquired specific drug resistance (resistant). In parallel, cells were cultured upon similar conditions in the absence of drug (parental). Proliferation of stable resistance was tested by MTT assay after 2-weeks of drug-free culture. Multi-drug resistance phenotype was ruled out by real-time PCR for MDR1 and MDR2-3 genes (Fig. S1) using published primers (17). As previously reported for a different resistant model (12), biological replicates were created by splitting the resistant cells after one month from resistance development, keeping them separate for additional 6 months before performing further experiments. Cells were periodically tested for their identity by short tandem repeat (STR) DNA profiling (18) and for Mycoplasma negativity (12).

### Treatments

Copanlisib, umbralisib, duvelisib, ibrutinib, zanubrutinib, acalabrutinib, everolimus, bimiralisib, vincristine, decitabine, lapatinib and LIN1632 were purchased from Selleckchem. Pirtobrutinib was purchased from MedChemExpress. Loncastuximab tesirine was kindly provided by ADC Therapeutics (Epalinges, Switzerland). Human recombinant HBEGF (CYT-119) was purchased from Prospec (Rehovot, Israel).

Response to single or drug combination treatments was assessed upon 72hr of exposure to increasing doses of drug followed by MTT assay. Sensitivity to single drug treatments was evaluated by IC50 (4-parameters calculation upon log-scaled doses) and area under the curve (PharmacoGX R package (19)) calculations. The beneficial effect of the combinations versus the single agents was considered both as synergism according to the Chou-Talalay combination index (20) and as potency and efficacy according to the MuSyC algorithm (21). For conditioned medium experiments, parental cells were cultured with 48h-conditioned medium from idelalisib-resistant, washed out in PBS and underwent MTT proliferation assay. Stimulation experiments were performed by adding recombinant HBEGF or PBS to the culture medium at final concentration of 30ng/mL at the moment of cell seeding.

### Genomics and data mining

See Supplementary materials.

### ELISA

For ELISA assays the conditioned medium was cultured for 72hr and collected. Then medium was filtered twice (22μm) and centrifuged at 4,000 rpm for 30 minutes in Amicon Ultra-4 tubes (Ultracel 3k, Merk Millipore) to remove cells and particles, and analyzed via ELISA. ELISA (Human HB-EGF Quantikine ELISA Kit, R&D Systems) was performed according to the manufacturer’s protocols. ELISA assays on frozen human serum samples were performed using Luminex Assay (R&D Systems) according to the manufacturer’s protocols. The serum samples were collected from patients treated with idelalisib or ibrutinib and enrolled on tissue banking protocols at Dana-Farber Cancer Institute; all patients signed written informed consent prior to a sample being drawn, and the protocols were approved by the Dana-Farber Harvard Cancer Center Institutional Review Board.

### Flow Cytometry (FACS) and protein analyses

Levels of p-AKT, p-BTK, p-PLCG2, p-mTOR and p-ERK were determined as previously described (22) (Table S1). Surface expression of ERBB4 (Table S2) was measured as previously described (23). For intracellular HBEGF expression levels, cells were first stained for surface CD19 (CD19-PE), then fixed and permeabilized, and finally stained for intracellular HBEGF (HBEGF-APC) (Table S2). Western blotting was performed to determine the expression of AKT/p-AKT, ERK/p-ERK and GAPDH (Table S3). Protein extraction, separation, and immunoblotting were performed as previously described (24).

### Immunofluorescence and confocal microscopy

Cells were permeabilized with PBS + 0.1% Triton X-100 10 minutes at room RT. To block unspecific staining samples were blocked for 1h with PBS + 5% BSA at RT before staining. Antibodies were diluted in PBS + 5% BSA. The primary antibodies used included rabbit monoclonal anti human p-AKT Ser473 (1:1000, Cell signaling), NF-κB/p65 (E379) C terminal (1:200, Abcam) and a mouse monoclonal anti human NF-κB/p65 (F-6) N terminal (1:100; SantaCruz) (Table S3). Samples were incubated overnight at 4°C. For immunofluorescence, the following secondary antibodies were used: goat anti-rabbit IgG labelled with Alexa 488 and donkey anti-mouse labelled with Alexa 555 (1:1000, Invitrogen) 1h at room temperature in the dark. Slides were counterstained after 3 washes of PBS with 0.3 μg/mL 4,6-diamidino-2-phenylindole (Sigma-Aldrich). Images including Z-stacks were acquired on a Leica SP8 gSTED with an objective with x63 magnification. Nuclear localization of p65 was quantified by ImageJ software.

### Pharmacological combination screening

Pharmacological screening was performed by exposing resistant cells in parallel with the parental counterpart, as a single or in combination with idelalisib, to 348 compounds from a custom library containing agents belonging to the following classes: “Kinase Inhibitory”, “Epigenetic Compound”, “PI3K/Akt Inhibitor”, “Apoptosis”, “Anti-cancer Compound”, and “MAPK Inhibitor” (Selleckchem, TX, USA), as previously described (12)x. All compounds, including idelalisib, were used at 1μM.

### Gene silencing and miRNA mimics

Small interfering RNAs were used for gene expression silencing. The control siRNA pool, human ERBB4 siRNA pool and human ERBB2 siRNA pool were purchased (Dharmacon GE Healthcare, now Horizon Discovery Ltd.). miRNA mimics, control, hsa-let-7c-5p, hsa-miR-29c-3p and hsa-miR-30b-3p were purchased (Qiagen). Karpas1718 cells (1 million per sample) were transfected with siRNA pools (200 pmol), or 100 pmols of each pool when silencing both ERBB4 and ERBB2, or with miRNA mimics (200 pmol) using 4D Nucleofector (Amaxa-Lonza Basel, Switzerland), with protocol CM-137, according to manufacturer instructions, and incubated for 48h to check RNA downregulation and 72h to check effect on proliferation.

## Results

### A MZL model of secondary resistance to the PI3K and BTK inhibitors

After nine months of exposure to the PI3Kδ inhibitor idelalisib, a stable resistant cell line (RES), bearing a 30-fold IC50 increase versus the parental cells, was obtained from the MZL cell line Karpas1718 (Fig. 1A). The resistance was confirmed to be stable repeating the IC50 after three weeks in medium containing no drug. Before proceeding with further characterization, multidrug resistance was ruled out by the demonstration of no increased expression levels of MDR1/2 by semiquantitative real-time PCR (Fig S1).

**Figure 1.**
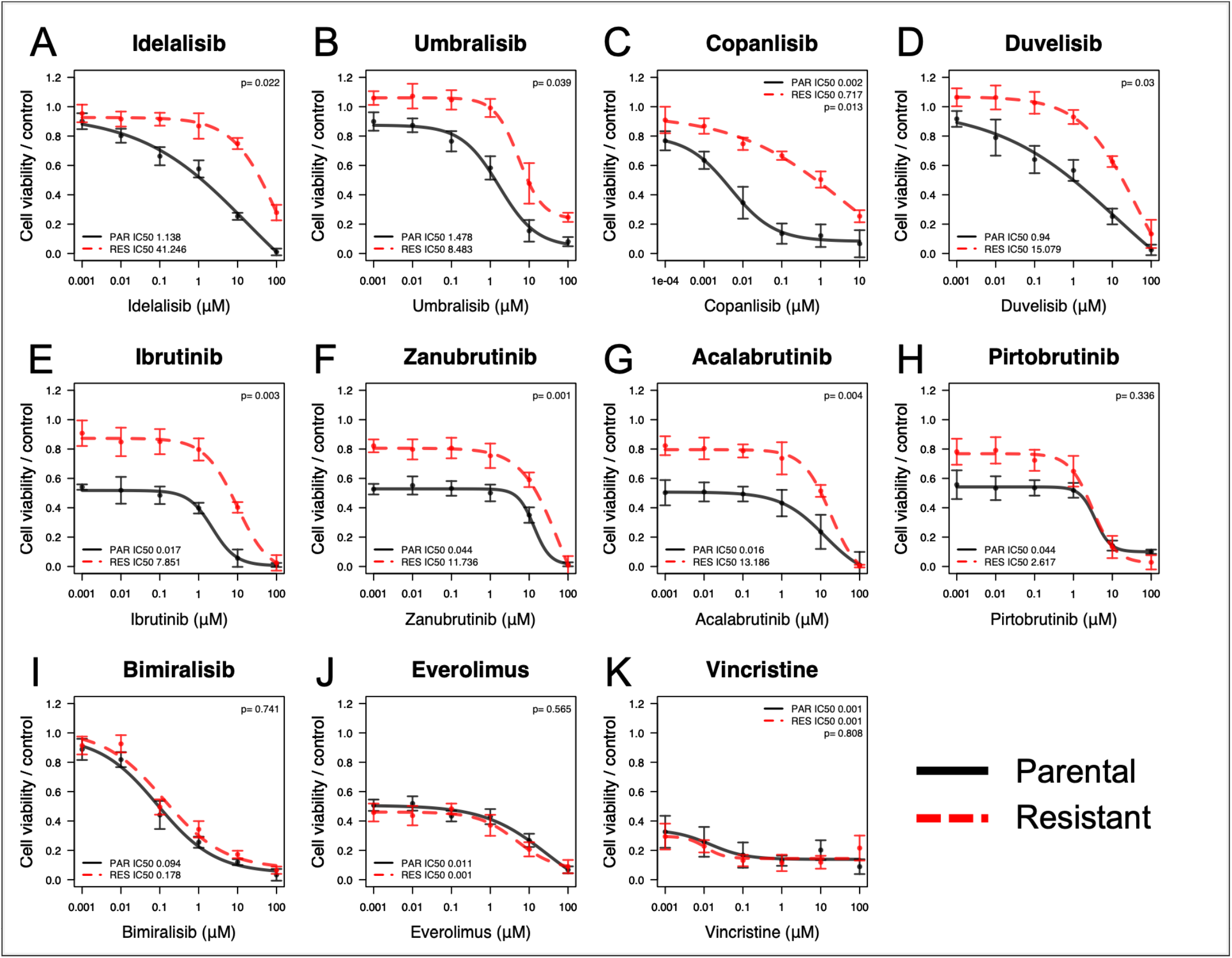
Profiles of drug sensitivity differ between parental and resistant lines. Acquired resistance was tested by MTT assay (72hr) in parental (black) and resistant (red dashed) cells of Karpas1718 line. Drug sensitivity was evaluated in resistant and parental cells for the PI3K inhibitors idelalisib (A), umbralisib (B), copanlisib (C), duvelisib (D), the BTK inhibitors ibrutinib (E), zanubrutinib (F), acalabrutinib (G), pirtobrutinib (H) the dual PI3K-mTOR inhibitor bimiralisib (I), the mTOR inhibitor everolimus (J) and the chemotherapy agent and MDR substrate vincristine (K). Error bars correspond to standard deviation of the mean; μM for micro molar. Data derived from at least three independent experiments. P values from a moderated t-test, statistically significant for p< 0.05.

Idelalisib-resistant Karpas1718 cell line showed resistance also against other FDA approved PI3K inhibitors (the PI3Kα/PI3Kδ inhibitor copanlisib, the PI3Kδ/PI3Kγ inhibitor duvelisib, and the PI3Kδ/CK1ε inhibitor umbralisib, Fig. 1B-D), and to the covalent (ibrutinib, zanubrutinib, acalabrutinib) and non-covalent BTK inhibitors (pirtobrutinib) (Fig 1E-H). Resistant cells exhibited were also resistant to the combination of rituximab and idelalisib (Fig S2). Conversely, the cell line was still sensitive to other agents such as the mTOR inhibitor everolimus, the dual PI3K/mTOR inhibitor bimiralisib and the chemotherapy agent, and MDR substrate, vincristine (Fig 1I-K).

Despite nor exonic nonsynonymous variants nor DNA copy number variations were acquired in the resistant cells when compared to their parental counterparts, the resistant Karpas1718 cell line markedly differed from its parental model based on RNA-sequencing (total RNA, miRNA) and methylation profiling (Fig S3A-B)(Table S4 SNV-CNV).

### Resistance to PI3K and BTK inhibitors is driven by ERBB signaling

The Karpas1718 resistant cells presented higher expression of genes involved in ERBB signaling (*HBEGF, NRG2, ERBB4*), cell cycle (*E2F2, MYBL2, CDC25B*), cell proliferation (*PBK, MKI67, TCL1A*), DNA recombination (*RAG1, RAG2*, histones) and TGFB signaling (*SMAD1, SMAD6, WNT8B*) when compared versus the parental cells. Conversely, BCR/NF-κB pathway (*SIRPA, TNF, BATF*), cytokine signaling (*IL10, IL6R, CCL22*), phosphatases (*DUSP2, DUSP4*) and genes down in clinical splenic MZL samples (*PLK2, MET, CAV1, PTEN*) were among the down-regulated transcripts (Fig 2, Fig S4A). Consistent with transcript level analyses, gene-set enrichment analyses (GSEA) showed enrichment in resistant of B cell proliferation, cell cycle, DNA recombination, ERBB, oxidative phosphorylation and proteosome. Of note, gene signatures up-regulated in diffuse large B cell lymphoma by PI3K or BTK inhibitors (idelalisib, duvelisib, bimiralisib and ibrutinib) were enriched in the resistant cells. Conversely, parental cells exhibited enrichment of BCR-NF-κB, TLR-MYD88, cytokine signaling and gene signatures repressed in splenic MZL clinical specimens and genes down-regulated by BET (JQ1), PI3K (copanlisib, AZD8835), and BTK (ibrutinib) inhibitors (Fig S4B).

**Figure 2.**
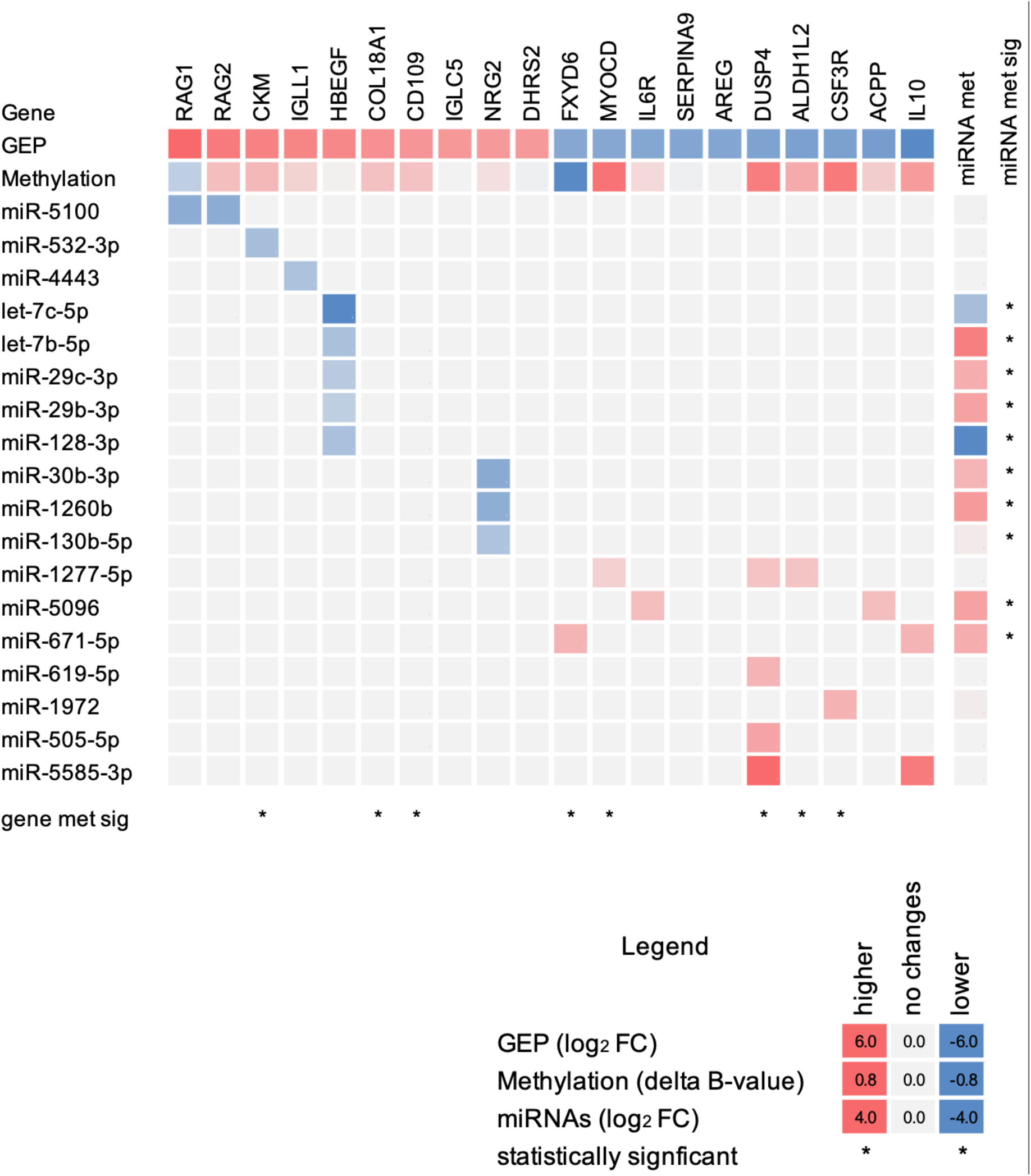
Multi-omics signature of resistant identifies activation of HBEGF-ERBB axis. Heatmap of RNA (GEP), methylation and miRNA profiles of resistant compared to parental. Heatmap values represent the differences between resistant and parental: fold change (log2 for RNA and miRNA) or delta Beta-value (methylation), red for enrichment in resistant and blue for parental. Columns correspond to gene expression (GEP, RNA-seq) and methylation (MethylationEPIC BeadChip, Illumina) profiles of the top-10 up-regulated and top-10 down-regulated genes; rows represent the differently expressed miRNAs (RNA-seq) with values in the column corresponding to the targeted gene. * for statistically significant differences (moderated t-test).

In the absence of mutations or DNA copy number changes we looked for changes at methylation and miRNA level. That might sustain the transcriptome changes. Comprehensive analysis of transcriptome, methylation and microRNA patterns identified miRNAs that were fully methylated in their promoters and down-regulated in resistant compared to parental cells. The negative regulators of the ERBB signaling miR-29b/c and let-7 targeting HBEGF (25), miR-30b and miR-1260 targeting NRG2 (predicted by TargetScan) and miR-625 and miR-3607 targeting ERBB4 (TargetScan) were among these (Fig 2, Table S5). We detected methylation in the promoter of *DUSP4* that may contribute to the down-regulation of this phosphatase in the resistant, which showed increased p-ERK (Fig 2, Fig S5).

The observed upregulation of genes involved in ERBB signaling activation was validated by demonstrating the over-expression of the ERBB4 on surface membrane by flow-cytometry (Fig 3A, S6) and the secretion of HBEGF into the medium by ELISA (Fig 3B). Consistent with autocrine secretion of HBEGF in resistant cells, intracellular staining of HBEGF was observed in resistant but not in parental (p<0.05, Fig 3C).

**Figure 3.**
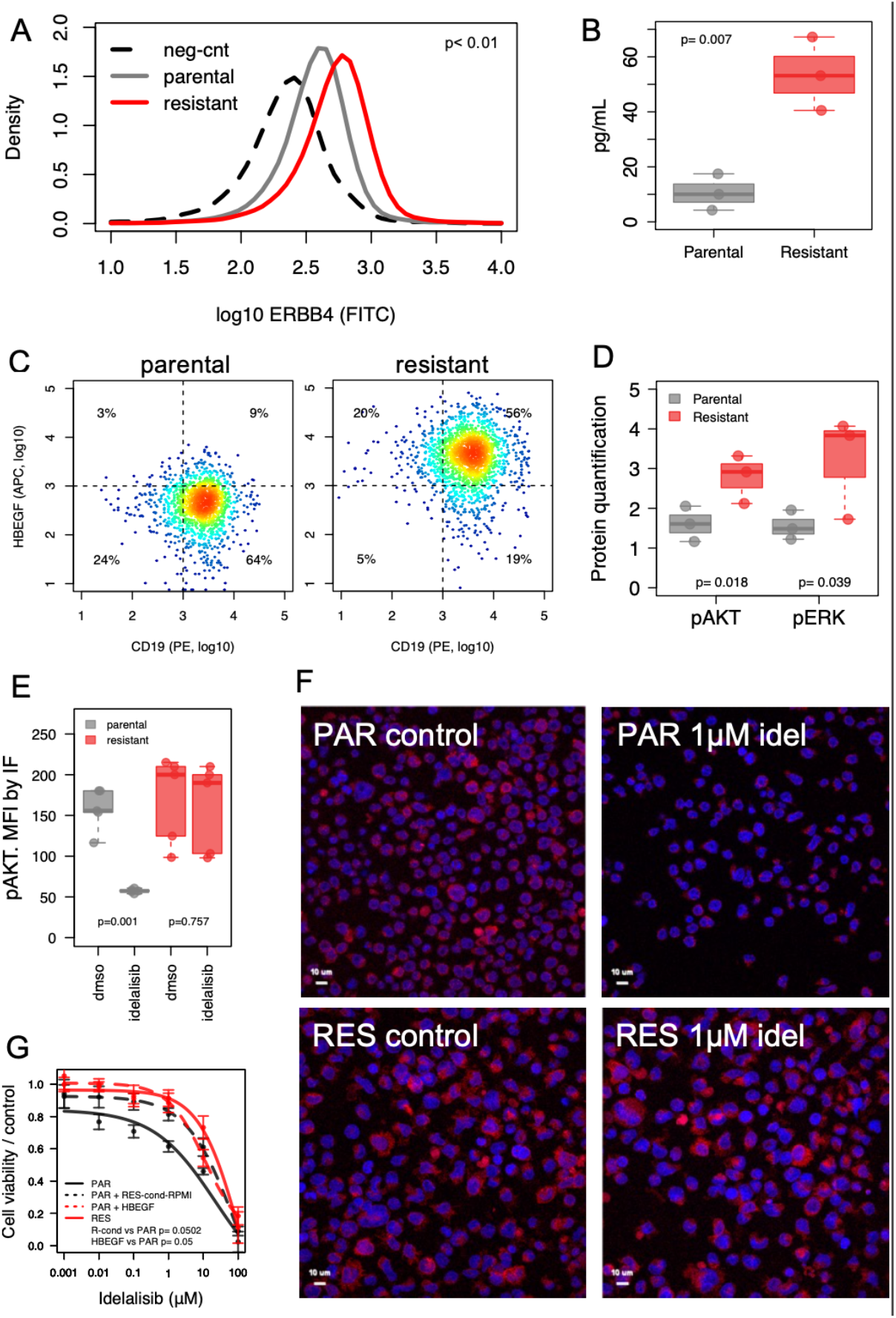
Proteome and secretome profiles of resistant differ from parental. (A) Expression of surface ERBB4 by FACS in parental (grey) and resistant (red) lines. Dashed black line for negative control (neg-cnt). Data derived from two independent experiments. (B) Secretion of HBEGF in parental (grey) and resistant (red) by ELISA. Data derived from three independent experiments. (C) Expression levels of surface CD19 (x-axis, PE-conjugated) and intracellular HBEGF (y-axis, APC-conjugated) by FACS in parental (left) and resistant (right) cells, percentage correspond to percentages of total viable cells. Data representative of two independent experiments. (D) Levels of protein phosphorylation by immunoblot in parental (grey) and resistant (red) cells. Values correspond to average of protein quantification normalized to GAPDH levels in three independent experiments. Error bars represent standard error of the mean. (E) Quantification of AKT phosphorylation by immunofluorescence, error bars represent standard error of the mean. Data derived from at least three independent experiments. P values from t-test, statistically significant for p< 0.05. (E) Detail of p-AKT expression by immunofluorescence in parental (PAR, top) and resistant (RES, bottom) upon DMSO (control, left) or 1μM idelalisib (right). (G) Cell viability by MTT assay of parental cells (PAR, black line) exposed to idelalisib in presence or not of conditioned medium from Karpas1718 idelalisib-resistant (RES, red line), cultured for 48h, PAR + RES-cond RPMI, dashed black line) or stimulated with 30ng/ml of recombinant HBEGF (dashed red line). Data derived from the average of three independent experiments. P values from a moderated t-test, statistically significant for p< 0.05.

Levels of p-AKT and p-ERK were also increased in resistant (immunoblotting, Fig 3D, Fig S7), and exposure to 1 μM of idelalisib decreased p-AKT and p-ERK in parental but had no effect in resistant cells, as observed by immunofluorescence and phospho-flow cytometry (p<0.05, Fig 3E-F, Fig S8A). Nuclear phospho-RELA was accordingly reduced in parental but not in resistant upon treatment with PI3Kδ inhibitors (p<0.05, Fig S8B). Moreover, culture of parental cells in conditioned medium from the resistant cells or stimulation with recombinant HBEGF reduced the sensitivity of the cells to idelalisib (Fig 3G).

Genetic silencing of *ERBB4* in resistant cells partially recover sensitivity not only to idelalisib but also to other PI3Kδ inhibitors, duvelisib, copanlisib or umbralisib. Nevertheless, knockdown of *ERBB2* had very limited effect in the sensitivity to PI3Kδ inhibitors, and concomitant silencing of both *ERBB4* and *ERBB2* gave no advantage in terms of sensitivity compared to ERBB4 single silencing, which significantly improved sensitivity to all PI3Kδ inhibitors but copanlisib (p=0.052) when compared to control (Fig 4A, Fig S9).

**Figure 4.**
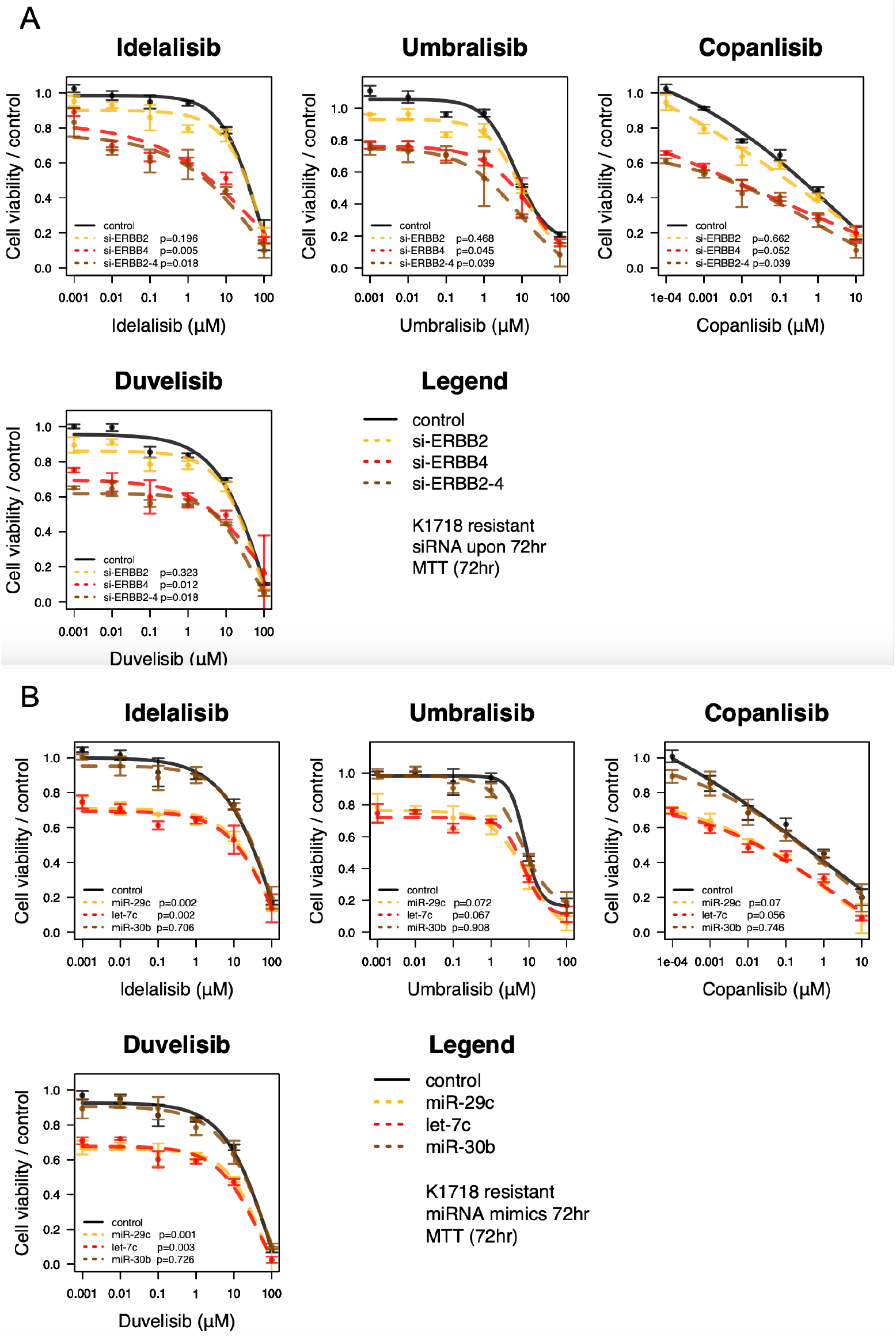
Genetic interfering with ERBB4, miR-29 or let-7 overcome resistance. (A) Small interfering RNAs were used for gene expression silencing of ERBB2 alone (yellow dashed line), ERBB4 alone (red dashed line) or concomitant silencing of ERBB2 and ERBB4 (brown dashed line). Black lines for resistant control. (B) miRNAs mimics were used for miR-29c expression (yellow dashed line), let-7c (red dashed line) or miR-30c (brown dashed line). Black lines for resistant control. Drug sensitivity was evaluated by MTT assay in resistant Karpas1718 cells for the PI3K inhibitors idelalisib, umbralisib, copanlisib and duvelisib. Data correspond of two independent experiments, error bars represent standard deviation of the mean. P values from a moderated t-test, statistically significant for p< 0.05. (C) Levels of HBEGF secretion by ELISA in resistant cells upon miRNA mimics for miR-29c (yellow), let-7c (red) or control (grey). Data derived from three independent experiments. P values from a moderated t-test, statistically significant for p< 0.05.

To further investigate the regulation of ERBB signaling in resistant cells due to the loss of some miRNAs, we evaluated changes on sensitivity to PI3Kδ inhibitors upon exposure to mimics of miRNAs targeting either *HBEGF* (let-7c and miR-29c) or *NRG2* (miR-30b). Supporting the role for HBEGF in the resistance to PI3Kδ inhibitors, the presence of let-7c or miR-29c, but not miR-30b, improved the sensitivity to PI3Kδ inhibitors (p<0.05, Fig 4B) and decreased HBEGF secreted levels (p<0.05, Fig 4C) in resistant cells.

### The addition of pan-ERBB inhibitor lapatinib reverts the resistance to PI3K inhibitors also in mantle cell lymphoma models

Since the resistance appeared due to the activation of the ERBB, we assessed whether the pharmacological inhibition of the pathway could be of benefit. We combined idelalisib with the pan ERBB inhibitor lapatinib in presence or not of recombinant HBEGF. Of note, either at baseline or upon HBEGF stimulation, addition of lapatinib (1μM) improved sensitivity to idelalisib (p<0.05, Fig 5A-B) and was synergistic in resistant and in the parental cells too (Fig S10). Since the deregulation observed in the let-7 family of miRNAs we wondered whether the addition of the LIN28 inhibitor LIN1632 might be of benefit in resistant cells. However, the combination of idelalisib with LIN1632 was of no benefit either in the parental nor resistant Karpas1718 cells, which do not express the drug target (Fig S11).

**Figure 5.**
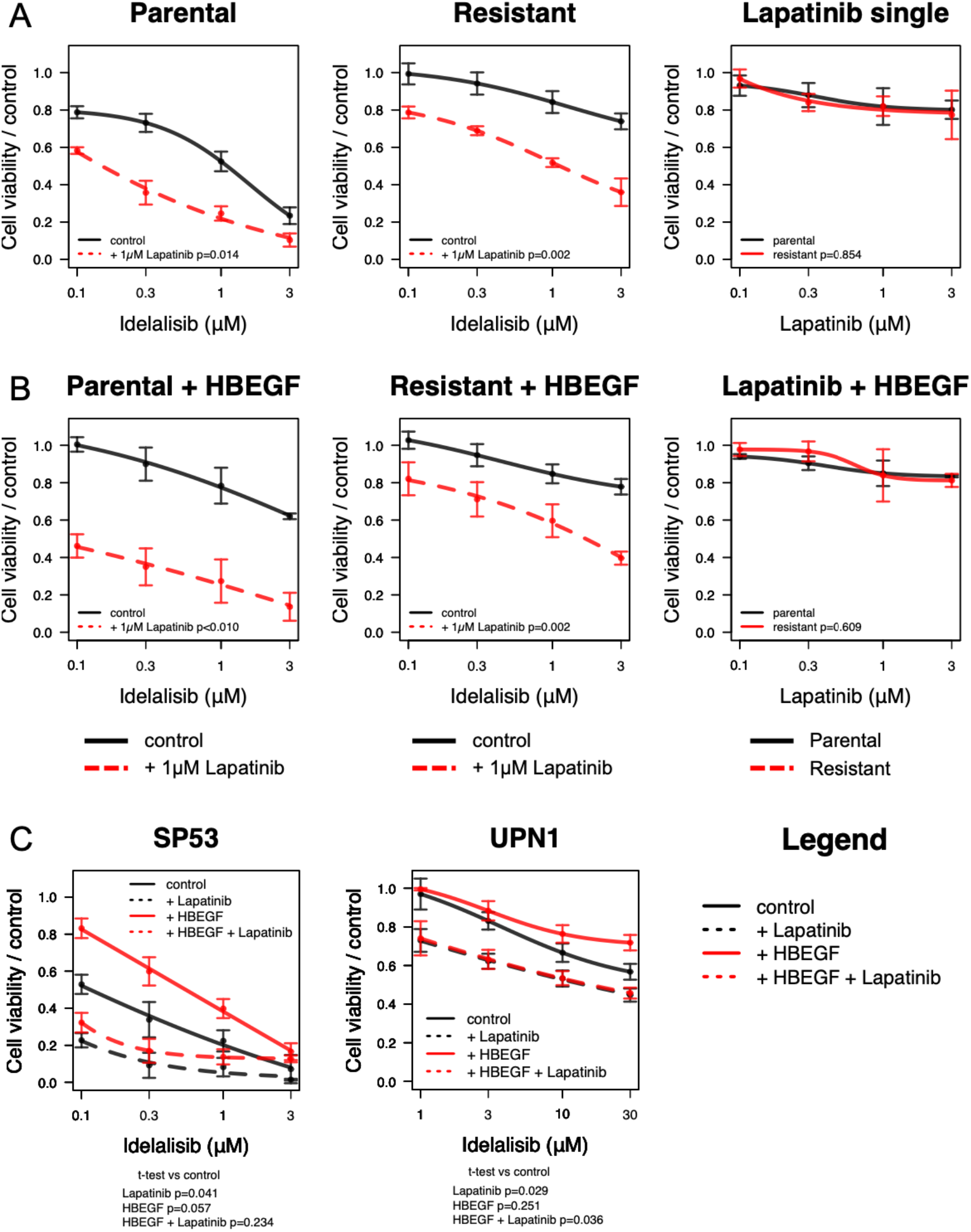
Pharmacological blocking of ERBB signaling recovers sensitivity to idelalisib. (A) Combination of idelalisib with 1μM of the ERBB inhibitor lapatinib increases sensitivity to idelalisib in either parental or resistant with no benefit as a single treatment. (B) Beneficial effect of the combination was observed even though upon stimulation with recombinant HBEGF. (C) Combination with ERBB inhibitors enhanced the potency of idelalisib in MCL models, either primary sensitive (SP53) or resistant (UPN1) to Idelalisib. Sensitivity to all treatments was tested by MTT assay upon 72h. Data correspond of three independent experiments; error bars represent standard deviation of the mean. P values from a moderated t-test, statistically significant for p< 0.05.

### The detected mechanisms of resistance are limited nor to PI3K inhibitors nor to the Karpas1718 model

We first observed that combination with ERBB inhibitors enhanced the potency of idelalisib also in two mantle cell lymphoma cell lines that were sensitive (SP53) or resistant (UPN1) to the PI3Kδ inhibitor, thus extending the significance of our findings beyond the Karpas1718 model (p<0.05, Fig 5C). Stimulation with HBEGF decreased sensitivity in the idelalisib-sensitive cell line (SP53) with limited effect in the idelalisib-resistant line UPN1 (Fig 5C).

We then took advantage of a large series of additional lymphoma cell lines we had previously characterized at transcriptome level and for their sensitivity to idelalisib (26). *HBEGF and ERBB4* expression levels were inversely correlated with idelalisib sensitivity, and the latter was positively correlated with *let-7c* and *miR-29c* levels also in these additional B cell lymphomas models (P<0.05, Fig S12) while not with other miRNAs of the same family let-7b and miR-29b.

The addition of recombinant HBEGF in the cell culture medium decreased the sensitivity not only to idelalisib but to other PI3K inhibitors, such as duvelisib, copanlisib and umbralisib (Fig 6A), and to the BTK inhibitors ibrutinib, zanubrutinib, acalabrutinib and pirtobrutinib (Fig 6B), while sensitivity to the dual PI3K-MTOR inhibitor bimiralisib was maintained (Fig 6A). The decrease in sensitivity to PI3K inhibitors was confirmed in the MZL models Karpas1718 (top panel) and VL51 (bottom panel, Fig 6A). Of interest, HBEGF stimulation affected the response to the BTK inhibitors as well, including the covalent-binding inhibitors ibrutinib, zanubrutinib, and acalabrutinib, and to the non-covalent BTK inhibitor pirtobrutinib in Karpas1718 parental, while resistant cells exhibited lower but still noticeable effect, in accordance with the constitutive autocrine secretion of HBEGF (Fig S6B). The decreasing on BTK inhibitors sensitivity upon HBEGF stimulation was observed in further *in vitro* lymphoma models too, including SP53 (MCL), OCILY10 and TMD8 (DLBCL) lines (Fig 6C). Similarly to the results obtained in the PI3K inhibitors, combination with the ERBB inhibitor lapatinib restored sensitivity to the BTK inhibitors in Karpas1718 lines and across additional B cell lymphoma models (dashed lines, Fig 6B-C). Altogether these results indicate that activation of ERBB signaling might drive resistance to PI3K and BTK inhibitors.

**Figure 6.**
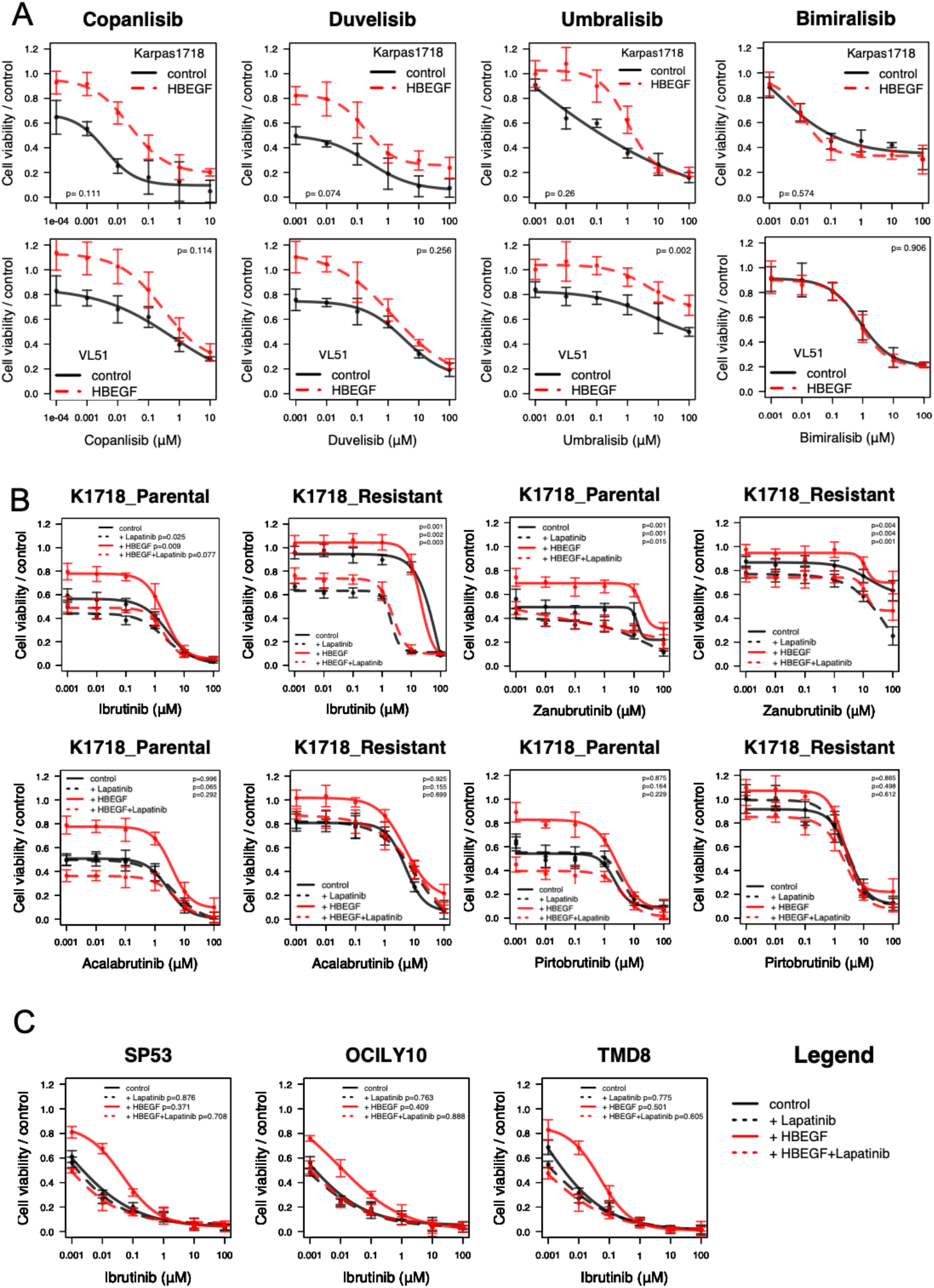
HBEGF stimulation confers resistance to other PI3K inhibitors and to BTK inhibitors. (A) Stimulation with HBEGF confer resistance to PI3K (A) or to BTK inhibitors (B). (A) Recombinant HBEGF (30 ng/mL, red line) decreased sensitivity to the PI3K inhibitors idelalisib, umbralisib, copanlisib and duvelisib in the two SMZL models Karpas1718 (top) and VL51 (bottom). (B) Recombinant HBEGF (30 ng/mL, red line) reduced sensitivity to the BTK inhibitors ibrutinib, zanubrutinib, acalbrutinib and pirtobrutinib in Karpas1718 parental and resistant lines. Sensitivity to the BTK inhibitors was restored by the addition of the ERBB inhibitor lapatinib (1μM, red dashed line). Control in black line and lapatinib single (1μM) in dashed black line. (C) Effect of HBEGF and/or lapatinib on the BTK inhibitor sensitivity across different B-cell lymphoma *in vitro* models: SP53 (MCL), OCILY10 and TMD8 (DLBCL). Sensitivity to all inhibitors was tested by MTT assay upon 72h. Data correspond of at least two independent experiments; error bars represent standard deviation of the mean. P values from a moderated t-test, statistically significant for p< 0.05.

### Pretreatment with epigenetic compounds recovers sensitivity to PI3K and BTK inhibitors

Among the 348 anti-cancer compounds tested, resistant cells showed acquired sensitivity to different epigenetic drugs, including DNA and histone methyltransferase inhibitors (Fig S13). Based on these results, as well the changes observed at methylation profiling, we tested a pretreatment with the demethylating agent decitabine given at low dose (100nM, 72hr), which improved sensitivity to idelalisib in resistant but not in parental cells. On the converse, neither single treatment nor concomitant combination of decitabine with idelalisib was beneficial in parental or resistant (Fig S14).

The response to ibrutinib also benefited from a pretreatment with the demethylating agent decitabine (100 nM, 72hr) or with the EZH2 inhibitor tazemetostat (5μM, 72hr) in resistant but not in parental cells, while single treatments or concomitant combination exhibited limited, or none advantage in all lines (Fig S15).

### The factors associated with resistance to PI3K and BTK inhibitors in cell lines are also observed in clinical specimens

To extend our findings to the clinical context, we investigated the expression levels for all factors associated with idelalisib-resistance in two series of MZL and in a large series of diffuse large B cell lymphoma (27–29). *HBEGF* and *ERBB4* were expressed in clinical specimens (Fig S16A-D).

We then took advantage of a previously reported gene expression profile of splenic MZL clinical specimens (28), determining the top 200 genes positively correlated (Pearson correlation) with *HBEGF* expression, and defining an *HBEGF* signature. When we applied the latter to our resistant model, the *HBEGF* signature was enriched among the genes more expressed in the resistant than in the parental cells (Fig S16E), emphasizing the similarities between our model and the clinical setting.

Finally, we investigated the relevance of the identified factors in the context of resistance to the PI3Kδ inhibitor idelalisib and to the BTK inhibitor ibrutinib in chronic lymphocytic leukemia (CLL) patients. Secreted levels of HBEGF were evaluated in the serum of CLL patients treated with idelalisib, comparing patients with primary or acquired resistance to the PI3Kδ or BTK inhibitors to patients responding to the drugs and paired for similar clinical features. In agreement with our *in vitro* data, serum samples from resistant patients exhibited significantly higher levels of HBEGF compared to serum samples of responders (Fig 7A). Upon ibrutinib treatment, HBEGF levels in patients wild-type *BTK* and wild-type *PLCG2* exhibited a trend of increasing at the time of progression compared to pre-treatment, suggesting a role for HBEGF in the response to ibrutinib as well (Fig 7B).

**Figure 7.**
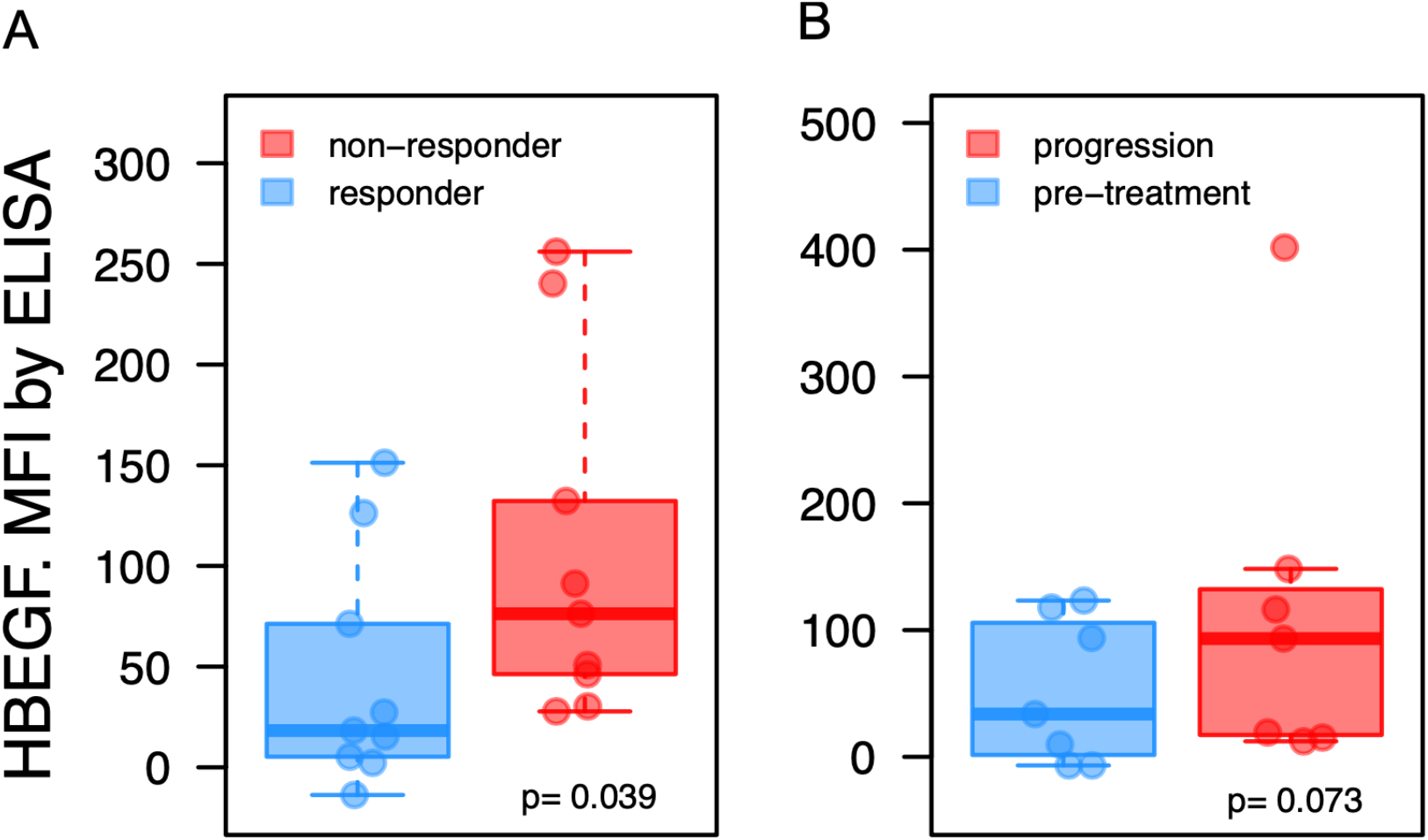
Higher HBEGF levels is detected in serum from idelalisib or ibrutinib resistant CLL patients. (A) HBEGF secretion was evaluated in the serum of CLL samples at the end of the treatment by ELISA (Luminex, R&D Systems). (A) Patients responding (responders, blue) or non-responding (non-responders, red) to idelalisib were compared by t-test. (B) HBEGF secretion in the serum of patients treated with ibrutinib was compared at the time of progression (red) with baseline levels previously of ibrutinib treatment (pre-treatment, blue). P for adjusted p-value from a t-test. MFI: mean fluorescence intensity.

## Discussion

We developed and characterized a novel *in vitro* model of secondary resistance to BTK and PI3K inhibitors in B cell lymphoma. Our results indicate that: i) activation of ERBB signaling driven by secreted ligands and upregulation of receptors can sustain resistance to BTK and PI3K inhibitors; ii) the expression of the genes sustaining the resistance is due to epigenetic reprogramming; iii) the resistance can be overcome using epigenetic agents and also drugs not usually employed in lymphoma patients, such as ERBB inhibitors, which could be tested in novel clinical trials.

The newly developed model of resistance originated from a MZL cell line kept for nine months under the PI3Kδ inhibitor idelalisib. Importantly, the resistance was not limited to idelalisib, but it extended to the combination with rituximab, to second generation PI3Kδ inhibitors (zandelisib), molecules targeting other kinases alongside PI3Kδ (copanlisib, duvelisib, umbralisib), and BTK inhibitors. The latter included both covalent (ibrutinib, zanubrutinib, acalabrutinib) and non-covalent inhibitors (pirtobrutinib).

In accordance with our results, DLBCL preclinical models with acquired resistance to idelalisib exhibit decreased sensitivity to BTK inhibitors, and ibrutinib-resistant models (driven by *BTK* or *TNFAIP3* mutations) lose response to other downstream BCR inhibitors, including idelalisib (9,15). Hence, the biological mechanisms observed for idelalisib can be extended to additional agents and new therapeutic strategies should be developed to escape acquired resistance to downstream B cell receptor targeted agents.

Our findings are in line with the notion that secreted factors can represent resistance factors to PI3K inhibitors, as described by our group and others (12,30). The Karpas1718 resistant cells secreted heparin-binding EGF (HBEGF) in the medium, which led to the activation of the ERBB pathway that blocked the anti-tumor activity of PI3K and BTK inhibitors. ERBB4 (HER4) is one of the four epidermal growth factors receptors, and it was upregulated together with its ligands HBEGF and neuregulin-2 (NRG2) in the resistant cell line. Besides being produced by lymphoma cells themselves, HBEGF and NRG2 are secreted in the tumor microenvironment by macrophages and T cells, respectively (31–33). Supporting a role for ERBB4 activation in BTK-inhibitor resistance, mutations in ERBB4 have been reported in MCL patients with primary resistance to ibrutinib (34).

The mechanism of resistance appeared driven by extensive methylation changes. Promoter methylation changes largely sustained the resistance via downregulation of miRNAs (miR-29 and let-7). Binding ERBB4 by its ligands activates the pro-survival downstream signaling cascade involving PI3K-AKT, PLCG, RAS-MEK-ERK and STAT (35). ERBB4 is associated with tumor development and progression in solid cancers, including triple negative breast cancer (TNBC) (36) but also in ALK-negative anaplastic large cell lymphomas (37). In TNBC, HBEGF is constitutively activated (38) whilst miR-29c, down-regulated in our model, is repressed (39) suggesting a role for the miR-29-HBEGF-ERBB4 axis in the activation of the PI3K-AKT signaling. Importantly, mir-29 directly targets HBEGF (25) and its role as tumor suppressor is reported in hematological malignancies including splenic MZL (40–42). Pretreatment with epigenetic compounds, such as demethylating agents and the EZH2 inhibitor tazemetostat, recovered sensitivity to PI3K and BTK inhibitors in our resistant model. In accordance with these results, treatment with demethylating agents increased the expression of miR-29 suggesting that levels of miR-29 might be modulated by methylation (25).

The role of miRNAs in causing resistance to idelalisib did not appear limited to miR-29. HBEGF is modulated by let-7 family of microRNAs (43). NRG2 is predicted to be regulated by miR-30 and miR-1260 microRNAs, both down-regulated in our model. MiR-1260 is associated with activation of ERBB and PI3K signaling cascades in solid cancers (44). DUSP4, downregulated and hypermethylated at the promoter level in the resistant cells, encodes a phosphatase that inactivates ERK and JNK MAP-kinases (45). DUSP4 is aberrantly methylated and repressed in DLBCL (46), and its silencing reduces the response to chemotherapy in HER2+ breast cancer cells (45). Therapeutically, the addition of the ERRB inhibitor lapatinib, FDA approved for patients with advanced or metastatic breast cancer (47), overcame resistance to idelalisib and it could be explored in the clinical setting for combinatorial regimens in the lymphoma setting for patients relapsing or progressing under PI3K and BTK inhibitors.

The mechanisms of resistance to BTK and PI3K inhibitors identified here might have been selected due to the background of the cell line, which reflect the genetic landscape of the recently described DMT SMZL subgroup, driven by mutations in genes involved in DNA-damage response, MAPK and TLR (48). Karpas1718 cells harbors lesions affecting cell proliferation, including *TP53* and *MYC*, and TLR signaling (*MYD88*). Thus, resistant cells of Karpas1718 model exhibited activation of ERBB signaling leading to increased cell proliferation, also in accordance with the proliferative genetic background of the cell line.

Importantly, the factors associated with resistance in the cellular model were validated across a large series of additional cell lines derived from different lymphoma types and their expression was shown in clinical specimens, including in the serum of idelalisib resistant and ibrutinib-treated and relapsed patients.

The enrichment of *RAG1* and *RAG2* genes among the transcripts upregulated in the resistant cells is in line with the previous observations that PI3Kδ and AKT inhibition increase the activity of these proteins, which are instead suppressed by PI3K and AKT signaling (49). In accordance as well with previous work on resistance to PI3K inhibitors, we observed activation of p-ERK (11,12), down-modulation of PTEN and activation of p-AKT (9) in the resistant cells.

In conclusion, the model of secondary resistance to the BTK and PI3K inhibitors, derived from the bona fide MZL Karpas1718 line has revealed novel ERBB4-driven mechanisms of resistance to the drugs and allowed the identification of a first series of active therapeutic approaches that can be further explored.

## Acknowledgments

This study was supported in part from the Swiss National Science Foundation (SNSF 31003A_163232/1) to EZ, DR and FB. JRB was supported by NIH RO1 CA 213442 (PI: Brown, Jennifer).

## Author contribution

AJA performed experiments, analyzed and interpret data, performed data mining, prepared the figures and cowrote the manuscript; SN performed silencing experiments and interpreted data; LC performed data mining; LB, GS, EC, EG, CT, AM, FS, performed experiments; AZ, FR, RBP, GS, VG performed flow-cytometry analyses; AR performed genomics experiments; MCM, ME performed methylation profiling experiments and data mining, SJ performed ELISA Luminex experiments; AS provided advice, JRB collected and characterized tumor samples; EZ and DR codesigned research, edited the manuscript, FB designed research, interpreted data, and cowrote the manuscript; all authors approved the final manuscript.

