## Supplementary material for "ERBB4-mediated signaling is a mediator of resistance to BTK and PI3K inhibitors in B cell lymphoid neoplasms"

*[1] Institute of Oncology Research, Faculty of Biomedical Sciences, USI, Bellinzona, Switzerland; [2] SIB Swiss Institute of Bioinformatics, Lausanne, Switzerland; [3] Centro di Riferimento Oncologico di Aviano – CRO, Aviano, Italy; [4] Josep Carreras Leukaemia Research Institute (IJC), Badalona, Barcelona, Catalonia, Spain; [5] Institute for Research in Biomedicine, Università della Svizzera italiana, Bellinzona, Switzerland; [6] Oncology Institute of Southern Switzerland, Bellinzona, Switzerland; [7] Faculty of Biomedical Sciences, USI, Bellinzona, Switzerland; [8] Chronic Lymphocytic Leukemia Center, Division of Medical Oncology, Dana-Farber Cancer Institute and Harvard Medical School, Boston, MA, USA; [9] Centro de Investigacion Biomedica en Red Cancer (CIBERONC), Madrid, Spain: [10] Institucio Catalana de Recerca i Estudis Avançats (ICREA), Barcelona, Catalonia, Spain: [11] Physiological Sciences Department, School of Medicine and Health Sciences, University of Barcelona (UB), Barcelona, Catalonia, Spain.*

*, equally contributed

### Supplementary materials and methods

Treatments

Response to single or drug combination treatments was assessed upon 72hr of exposure to increasing doses of drug followed by MTT assay. Cells were plated in 96-well plates at a concentration of 20,000 per well in non-phenol RPMI-1640 (Gibco Invitrogen, Basel, Switzerland) supplemented with 10% fetal bovine serum (Gibco) and 1% penicillin-streptomycin (Gibco). After 4 hours of seeding, cells were exposed treatments. Sensitivity to single drug treatments was evaluated by IC50 (4-parameters calculation upon log-scaled doses) and area under the curve (PharmacoGX R package (1)) calculations. The beneficial effect of the combinations versus the single agents was considered both as synergism according to the Chou-Talalay combination index, as previously described (2), and as potency and efficacy according to the MuSyC algorithm (3). For conditioned medium experiments, parental cells were cultured with 48h-conditioned medium from idelalisib-resistant, washed out in PBS and underwent MTT proliferation assay. Moderated t-test (*limma* package in R environment) was performed to determine statistically significant differences in drug response experiments ($p<0.05$).

Flow Cytometry (FACS) and protein analyses

Surface expression of ERBB4 and CD19 (Table S1) were measured as previously described (4). Levels of p-AKT, p-BTK, p-PLCG2, p-mTOR and p-ERK were determined as previously described (5) (Table S2). Levels of protein expression were analyzed by FACS with the flowCore R package (6).

Immunoblotting was performed to determine the expression of AKT/p-AKT, ERK/p-ERK JAK/p-JAK, STAT/p-STAT and GAPDH (Table S3). Protein extraction, separation, and immunoblotting were performed as previously described (2).

Genomics

For whole exome sequencing (WES), genomic DNA was enriched in protein coding sequences using the exome capture SureSelect XT library preparation (v6, 58 Mb; Agilent Technologies), according to the manufacturer's protocol. The captured targets were subjected to next generation sequencing using the HiSeq 2500 analyzer (Illumina) with the paired-end 2x125 bp read option, following the

manufacturer's instructions, to obtain a 14x coverage. Exome capture and next generation sequencing were performed at the HiSeq Service of Fasteris SA (Plan-les-Ouates, Switzerland).
Transcriptome sequencing (RNA-Seq) was done using the TruSeq RNA Sample Prep Kit v2 for Illumina (Illumina, San Diego, CA, USA) as previously described (7).
For small RNA-Seq, cDNA libraries were assembled using total RNA prepared using the SMARTer smRNA-Seq Kit for Illumina (Clontech Laboratories, Inc. USA). Briefly, the total RNA suspension was polyadenylated by the Poly(A) Polymerase at 16°C for 5 minutes on a thermal cycler, then cDNA synthesized using PrimeScript Reverse Transcription, primed by the 3' smRNA dT primer. cDNA was amplified and full-length Illumina adapters were added via PCR. PCR products were purified using the NucleoSpin Gel and PCR Clean-Up kit. Libraries were amplified using 10 cycles of PCR on thermocycler, then quantified on a Qubit 4 Fluorometer (Invitrogen, USA) and analyzed on an Agilent Bioanalyzer High Sensitivity DNA chip for qualitative control. The cDNA libraries were size selected to enrich for <150bp using Agencourt AMPure XP beads, first selection with a (0.8X) of beads and second selection with a (2.2X) of beads. Libraries were then sequenced on a NextSeq500 Illumina using 75 cycles.
Methylation profiling was done using the MethylationEPIC BeadChip Infinium following the manufacturer's instructions for the automated processing of arrays with a liquid handler (Illumina Infinium HD Methylation Assay Experienced User Card, Automated Protocol 15019521 v01), as previously described (8).

Data mining

For WES, quality control on raw reads was performed with FastQC (9). Paired-end reads were aligned to human reference sequence GRCh37 using the Burrows–Wheeler Aligner (BWA version 0.6.1) (10). Potential PCR duplicates were removed using SAM tools command (11). Mapping quality score recalibration and local realignment around insertions and deletions (indels) were performed using the Genome Analysis Toolkit (GATK) (12). Single nucleotide variants and small indels were called separately using the GATK Unified-Genotyper (13). Annovar tool (14) was used for functional annotation of variants, and all mutations found were manually checked and explored using the Integrative Genomic Viewer 2.03 (15). The list of putative acquired variants was identified using the following filtering constrains. Mutations present in the parental cell lines were excluded, as well as known germline variants reported at dbSNP137 and the 1000 Genomes Project (16). We considered of our interest exonic and nonsynonymous variants, including stop-gain single nucleotide variants, splicing, and frameshift indels. We excluded non-exonic variants and synonymous mutations. We also retained variants already reported in the COSMIC database (17). Since samples underwent both whole exome sequencing and RNA-sequencing, variants were also investigated for the correspondence between whole exome and RNA-sequencing. Finally, only variants transcribed to RNA were considered for further analyses.
For RNA-Seq, data were analyzed as previously described (7). Differentially expressed genes were calculated with moderated t-test on RNA-seq. The false discovery rate (FDR, Benjamin-Hochberg correction) was calculated to control for false positives. FDR <0.05 and absolute fold-change higher than 2 was considered significant. Functional analysis was performed on the collapsed gene symbol list using GSEA (Gene Set Enrichment Analysis) with the MSigDB (Molecular Signatures Database) C2-C7 gene sets (18,19), and SignatureDB database (https://lymphochip.nih.gov/signaturedb/). Statistical tests were performed using the R environment (R Studio console; RStudio, Boston, MA, USA).
For small RNA-Seq, we first carried out a pre-processing step with Cutadapt (20) to identify and remove all of non-biological part of the reads (adapters). Then, a quality control step was performed using FastQC where we collected key information about the quality of sequencing reads including quality score distribution along the reads, GC content, read length, and level of sequence duplication. Once the FASTQ files have been validated, BWA aligned the reads to the human reference genome (hg38). We then quantified the known microRNAs present in miRBase (21), counting the reads mapped to their loci with featureCounts (22). We set the -O flag to have reads that map to several overlapping microRNAs assigned to all of them. We also set the -s parameter to 1 to only count reads that map to

the same strand as the microRNA, and the -M flag to make sure we count multi mapping reads. We compensated for different sequencing depths using the TMM normalization (normalization step), and calculated the log2 counts-per million values (lcpm). Log count-per-million values were inputed to principal component analysis (PCA) or multi-dimensional scaling plot to get a global look of how similar the microRNA expression profiles are in the different samples. Limma identified the modulated miRNAs between the two phenotypes of interest.

For methylation profiling, the DNA methylation beta values were obtained from the raw IDAT files by using the minfi package in R. The data was normalized with the ssNoob method and positions with a detection P-value greater than 0.01, NoCG Start as well as probes with SNPs, multihit start probes and XY chromosome probes were removed from the analysis. The differentially methylated positions between groups with a Benjamini-Hochberg adjusted p-value lower than 0.05 and an absolute methylation difference greater than 0.3 were selected from a linear model calculated with the limma R package.

Finally, unsupervised multidimensional scaling plot were used to visualize resistant and parental multi-omics profiles.

Gene silencing

Small interfering RNAs were used for gene expression silencing. The control siRNA pool, human ERBB2 siRNA pool and human ERBB4 siRNA pool 200 pmol were purchased (Dharmacon GE Healthcare, Horizon Discovery Ltd., Cambridge, UK). Karpas1718 cells (1 million per sample) were transfected with siRNA pools (200 pmol), or 100 pmols of each pool when silencing both ERBB2 and ERBB4, using 4D Nucleofector (Amaxa-Lonza, Basel, Switzerland), with protocol CM-150, according to manufacturer instructions, and incubated for 48h to check RNA downregulation and 72h to check effect on proliferation.

Intracellular staining

cells were stained for surface CD19 (CD19-PE 1:100 dilution, Table S2), fixed with 4% formaldehyde PBS for 15 minutes at room temperature; and permeabilized in 0.5% v/v Tween-20 PBS for 10 minutes at room temperature. Cells were then stained for intracellular HBEGF (HBEGF-APC dilution 1:100, Table S2) in 0.1% Tween20 PBS, 30 minutes room temperature. Cells were washed twice (2mL 0.1% Tween20/Triton PBS, spin 1200rpm 5 minutes), resuspended on 500µL 1% FBS PBS and signal was measured by flow cytometry.

### Supplementary tables

**Table S1**. Panel of kinases analyzed in the Phospho Flow experiments of parental and resistant cells by flow cytometry.

| protein | fluorochrome | company | # catalog |
|---|---|---|---|
| ERK 1/2 (pT202/Y204) | Alexa488 | BD Biosciences | 612592 |
| AKT (pS473) | Alexa647 | BD Biosciences | 560343 |

**Table S2**. Panel of antibodies used in flow cytometry experiments.

| Protein | fluorochrome | company | # catalog |
|---|---|---|---|
| ERBB4 | FITC | Santa Cruz | 8050 |
| | 2nd ab | Dako | F0479 |
| ERBB4 | Alexa488 | R&D Systems | FAB11311G |
| CD19 | PE | BD Bioscience | 555413 |
| HBEGF | APC | R&D Systems | IC259A |

**Table S3**. Panel of proteins tested in the immunoblotting experiments of parental and resistant cells by western blotting.

| source | protein | company | # catalog |
|---|---|---|---|
| rabbit polyclonal | α-AKT | Cell Signaling | 9272 |
| rabbit polyclonal | α-p(S473) AKT | Cell Signaling | 4060 |
| rabbit polyclonal | α-ERK1/2 | Cell Signaling | 4696 |
| rabbit polyclonal | α-p(Y204) ERK | Santa Cruz | 7383 |
| mouse monoclonal | α-GAPDH | CNIO | FF26A/F9 |

**Table S4 (Excel file**). Whole exome sequencing data, including single nucleotide variants and copy number variants.

**Table S5 (Excel file).** Multi-omics analyses. Output tables of the moderated t-tests comparing transcriptome, microRNA and methylation profiles of resistant and parental. Output results of Gene Set Enrichment Analysis comparing GEP data of resistant and parental.

**Table S6 (Excel file).** Clinical information on the series of serum samples from CLL clinical patients exposed to idelalisib or to ibrutinib. Samples were longitudinally acquired at different time points. Patients were paired according to similar clinical features.

#### Supplementary Figures

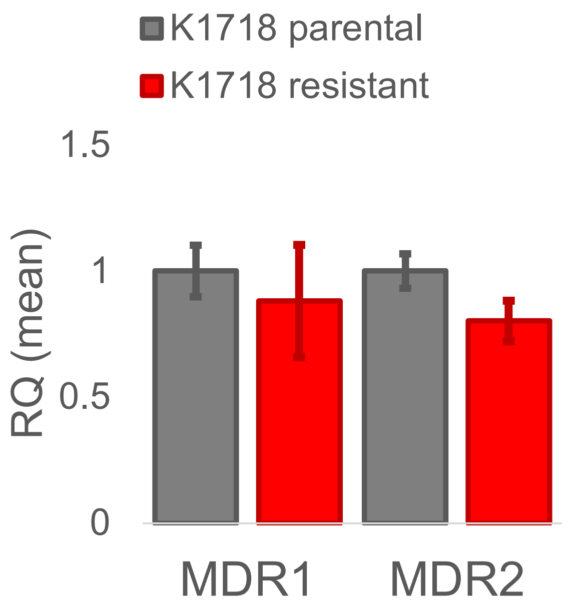


**Figure S1**. Multi-drug resistance phenotype was ruled out by the expression of MDR1 (left) and MDR2 (right) genes by real time PCR. RQ values calculated by the DDCt method. Karpas1718 (K1718) parental lines in grey and resistant in red. Data derived from two independent experiments; error bars represent standard deviation of the mean.

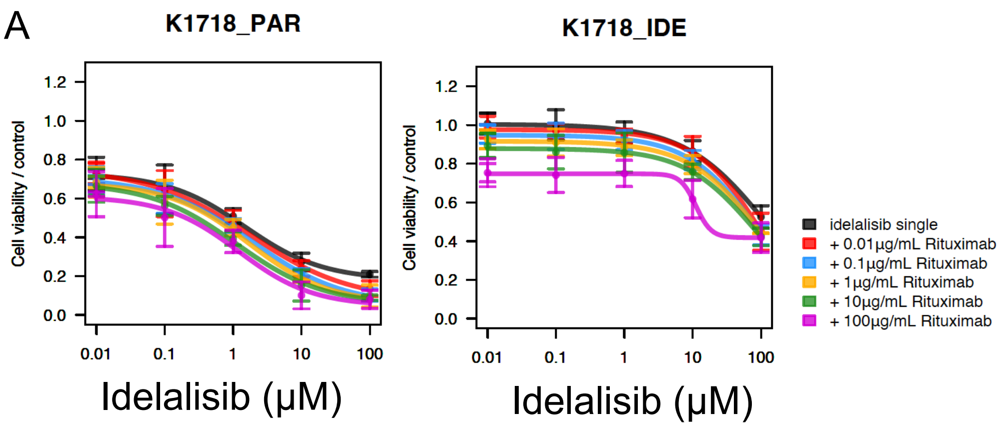


| p-value from moderated t-test comparing each combination to idelalisib single | | |
|---|---|---|
| Combination | Parental | Resistant |
| 0.01μg/mL Rituximab | 0.792 | 0.272 |
| 0.1μg/mL Rituximab | 0.605 | 0.053 |
| 1μg/mL Rituximab | 0.525 | 0.12 |
| 100μg/mL Rituximab | 0.42 | 0.002 |
| 10μg/mL Rituximab | 0.274 | <0.001 |

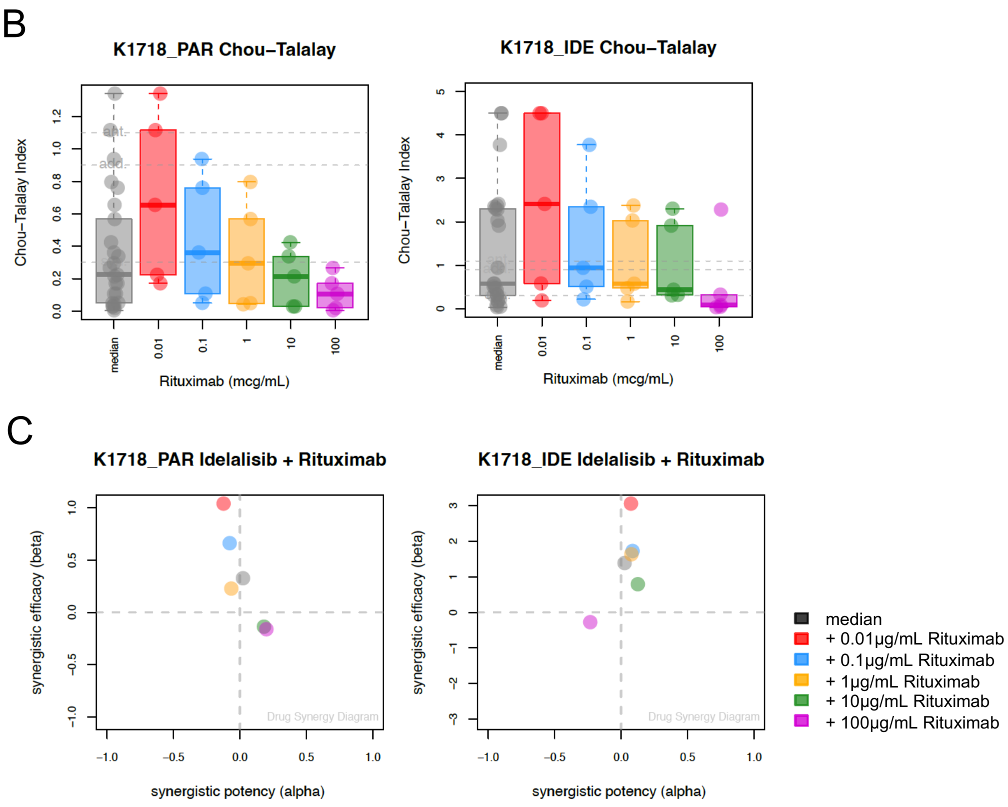


**Figure S2**. (A) Cell viability for the combination of idelalisib and the CD20 blocking antibody rituximab in parental and resistant by MTT assay (72h). Bars correspond to the mean of two independent experiments. Error bars represent standard deviation of the mean. Table contains p-values from a moderated t-test comparing each combination to idelalisib as single agent. The benefit of the combination was assessed both as synergism according to the Chou-Talalay combination index (B) (23) and as potency (x-axis) and efficacy (y-axis) according to the MuSyC algorithm (C) (3). K1718_PAR: parental, K1718_IDE: resistant.

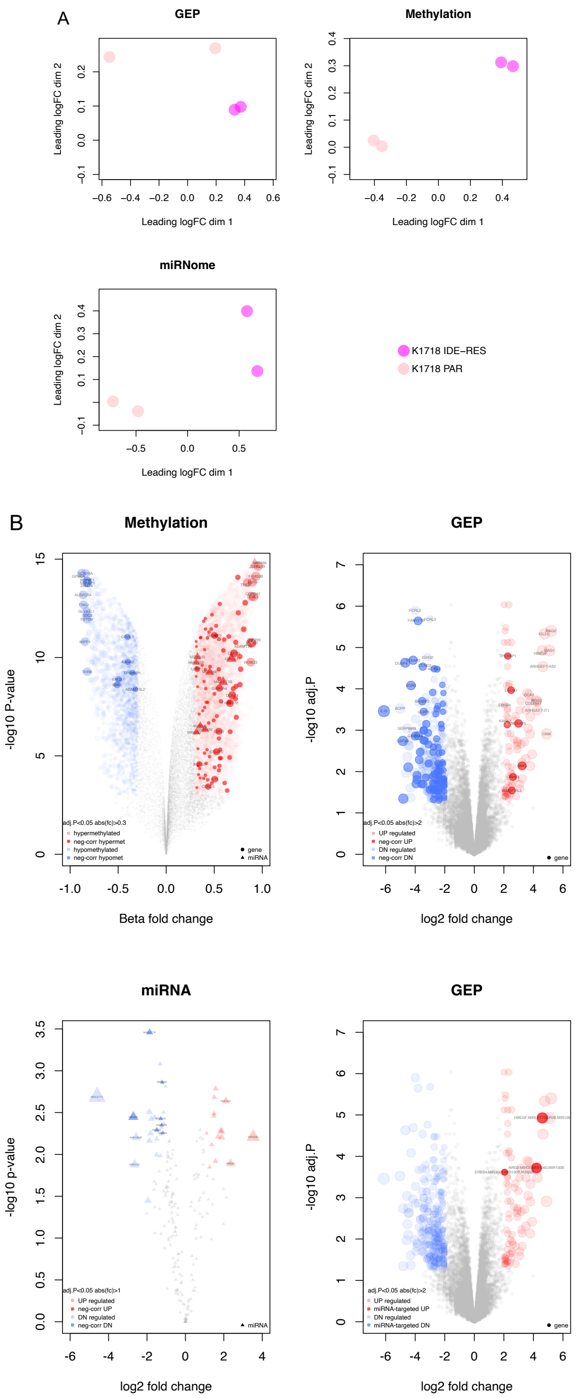


**Figure S3**. (A) Multidimensional scaling plot showing a 2-dimensional projection of distances across parental and resistant clones for RNA (GEP, top left), methylation (top right) and miRNA (bottom left) profiles. Magenta dots for resistant and pink for parental. (B) Vulcano plots on methylation (top left), gene expression (RNA-seq, top right) and microRNA (RNA-seq, bottom left) profiles of Karpas1718 resistant compared to parental. GEP-miRNA profiles integration. Moderated t-test (limma R Package) was performed comparing resistant to parental: delta Beta-value (methylation), fold change (RNA-seq), adj.P-value for Bonferroni correction of the nominal p-value. Dots represents genes and triangles represent miRNAs. Red corresponds to higher values in resistant, and blue higher values in parental. The genes or miRNAs inversely correlated with methylation are highlighted in darker color: hypomethylated and overexpressed in dark red (neg-corr UP) and hypermethylated and repressed in dark blue (neg-corr DN).

A

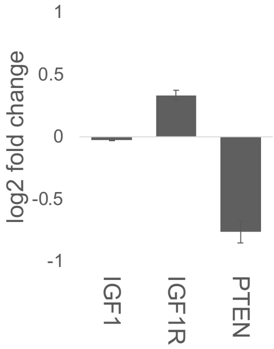


B

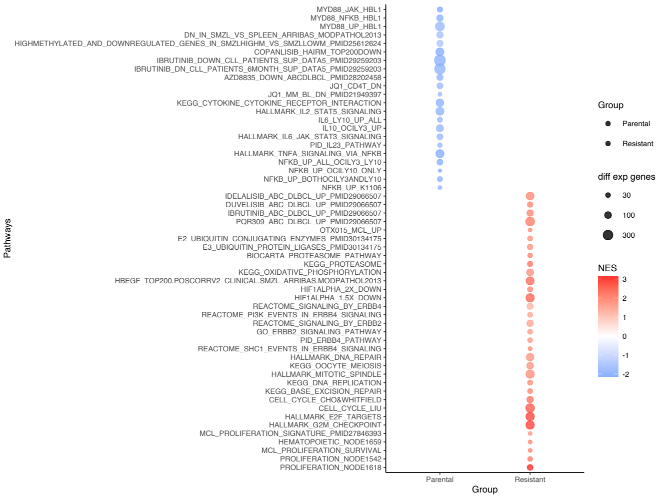


**Figure S4.** (D) RNA expression of IGF1, IGF1R and PTEN in parental and resistant of Karpas1718 lines. Bars represents log2-scaled fold change by RNA-seq, * for statistically significant differences. (B) Gene set enrichment analyses (GSEA, Broad Institute) comparing parental and resistant per each line. Statistically significant differences with values of absolute NES higher than1, P <0.05 and FDR <0.25.

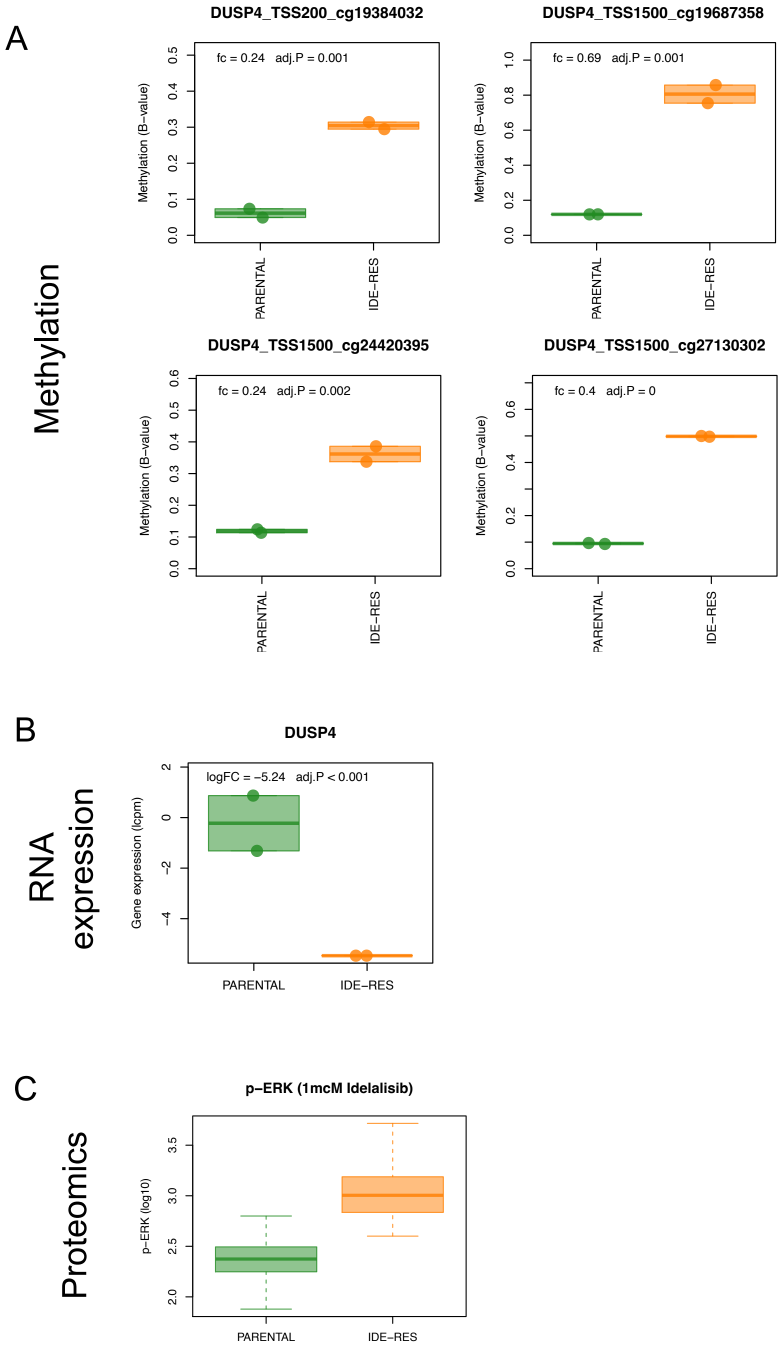


**Figure S5.** Parental (green) and resistant (orange) levels of DNA promoter methylation (A) and gene expression (RNA-seq, B) of DUSP4, and p-ERK upon 1μM of idelalisib (Flow cytometry).

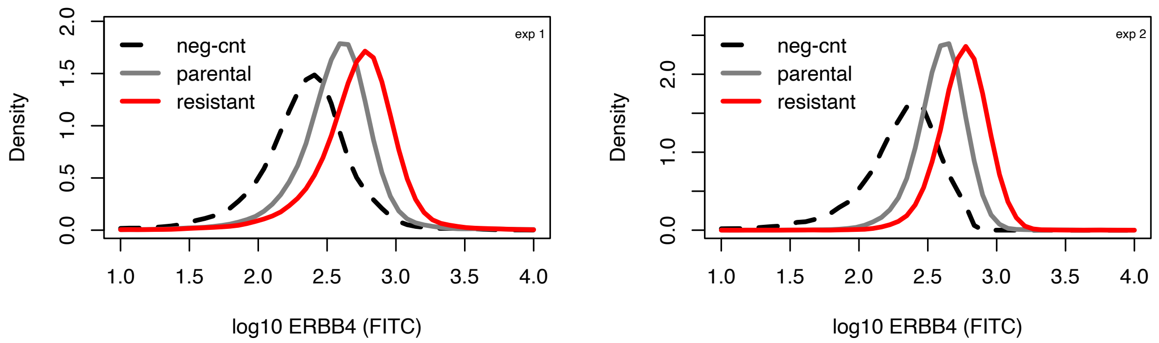


**Statistics (log MFI by FACS)**

Welch Two Sample t-test (exp 1+2)

1st exp p-val= 0 2nd exp

| 1st exp | 2nd exp |
|---|---|
| neg-cnt= 2.354 | neg-cnt= 2.334 |
| parental_replicate1= 2.593 | parental_replicate1= 2.624 |
| parental_replicate2= 2.551 | parental_replicate2= 2.574 |
| resistant_replicate1= 2.729 | resistant_replicate1= 2.778 |
| resistant_replicate2= 2.751 | resistant_replicate2= 2.748 |

**Statistics (log MFI by FACS)**

Welch Two Sample t-test (exp 1+2)

1st exp p-val< 0.01 2nd exp

| 1st exp | 2nd exp |
|---|---|
| neg-cnt= 2.354 | neg-cnt= 2.334 |
| parental_replicate1= 2.593 | parental_replicate1= 2.624 |
| parental_replicate2= 2.551 | parental_replicate2= 2.574 |
| resistant_replicate1= 2.729 | resistant_replicate1= 2.778 |
| resistant_replicate2= 2.751 | resistant_replicate2= 2.748 |

**Figure S6**. Expression levels of surface ERBB4 was measured by FACS in Karpas1718 parental and resistant. Two independent experiments were performed: exp1 (left panel) and exp2 (right panel). Density plots show the median MFI values of two replicates from each experiment: negative control (dotted black), parental (grey), resistant (red). Results are presented from Welch Two Sample test performed on MFI values of the two experiments.

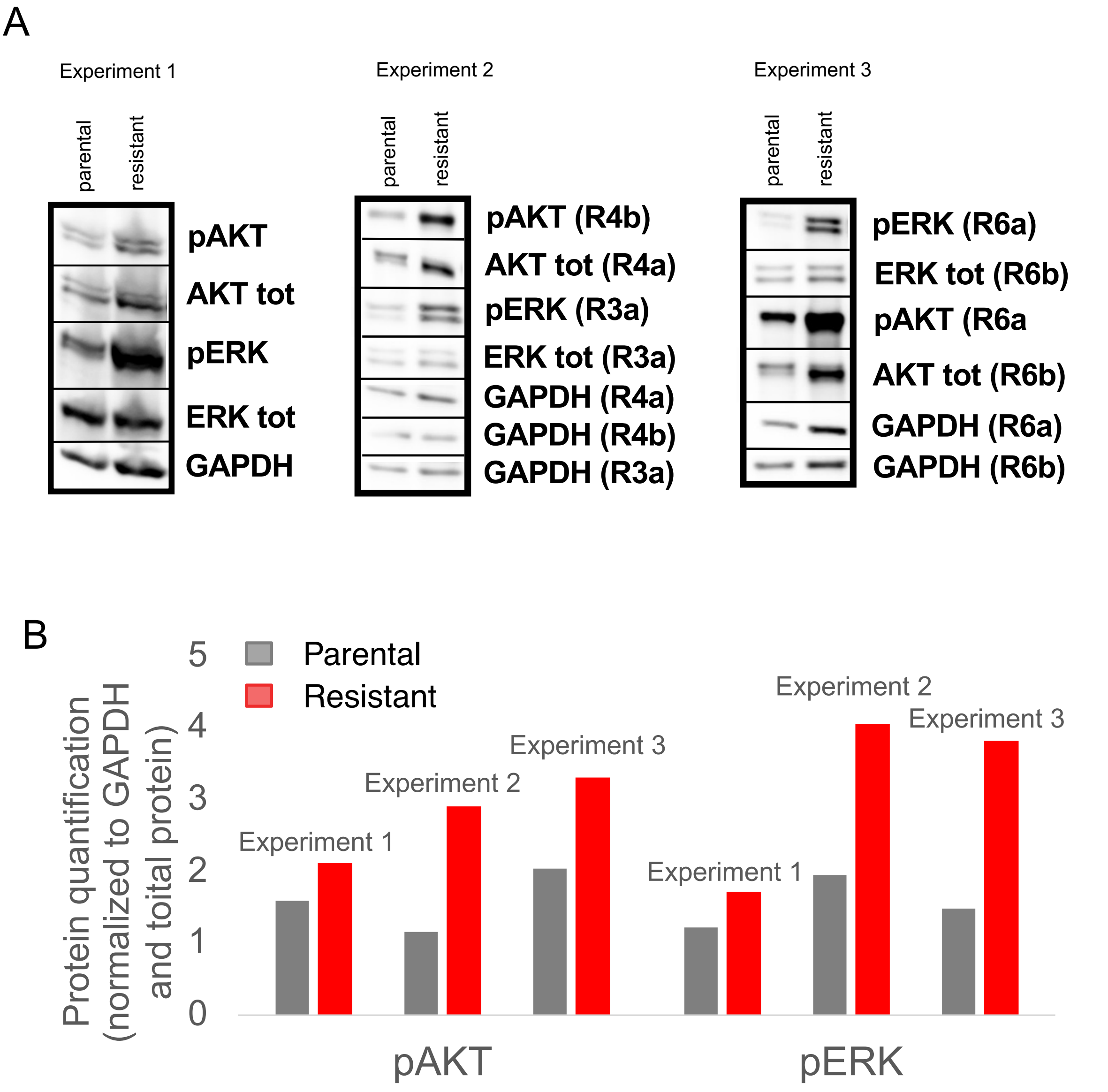


| protein | experiment 1 | | experiment 2 | | experiment 3 | | statistics | | | | |
|---|---|---|---|---|---|---|---|---|---|---|---|
| | protein quantification parental (normalized to GAPDH and total protein) | protein quantification resistant (normalized to GAPDH and total protein) | protein quantification parental (normalized to GAPDH and total protein) | protein quantification resistant (normalized to GAPDH and total protein) | protein quantification parental (normalized to GAPDH and total protein) | protein quantification resistant (normalized to GAPDH and total protein) | median parental | median resistant | std error parental | std error resistant | t-test, p-value parental vs resistant |
| pAKT | 1.606 | 2.119 | 1.163 | 2.915 | 2.053 | 3.317 | 1.606 | 2.915 | 0.121 | 0.166 | 0.018 |
| pERK | 1.222 | 1.725 | 1.955 | 4.068 | 1.486 | 3.835 | 1.486 | 3.835 | 0.101 | 0.351 | 0.039 |

**Figure S7.** (A) Immunoblotting was performed to measure protein levels in parental and resistant clones for pAKT/AKT and pERK/ERK in three independent experiments. GAPDH was used as a loading control. (B) Protein quantification was done normalizing first to GAPDH and then to total protein. Barplot represents the quantification in each of the independent experiments, parental in grey, resistant in red. Median and standard error of the mean for parental and resistant levels in the three experiments. Moderated t-test was performed to determine statistically significance ($p<0.05$).

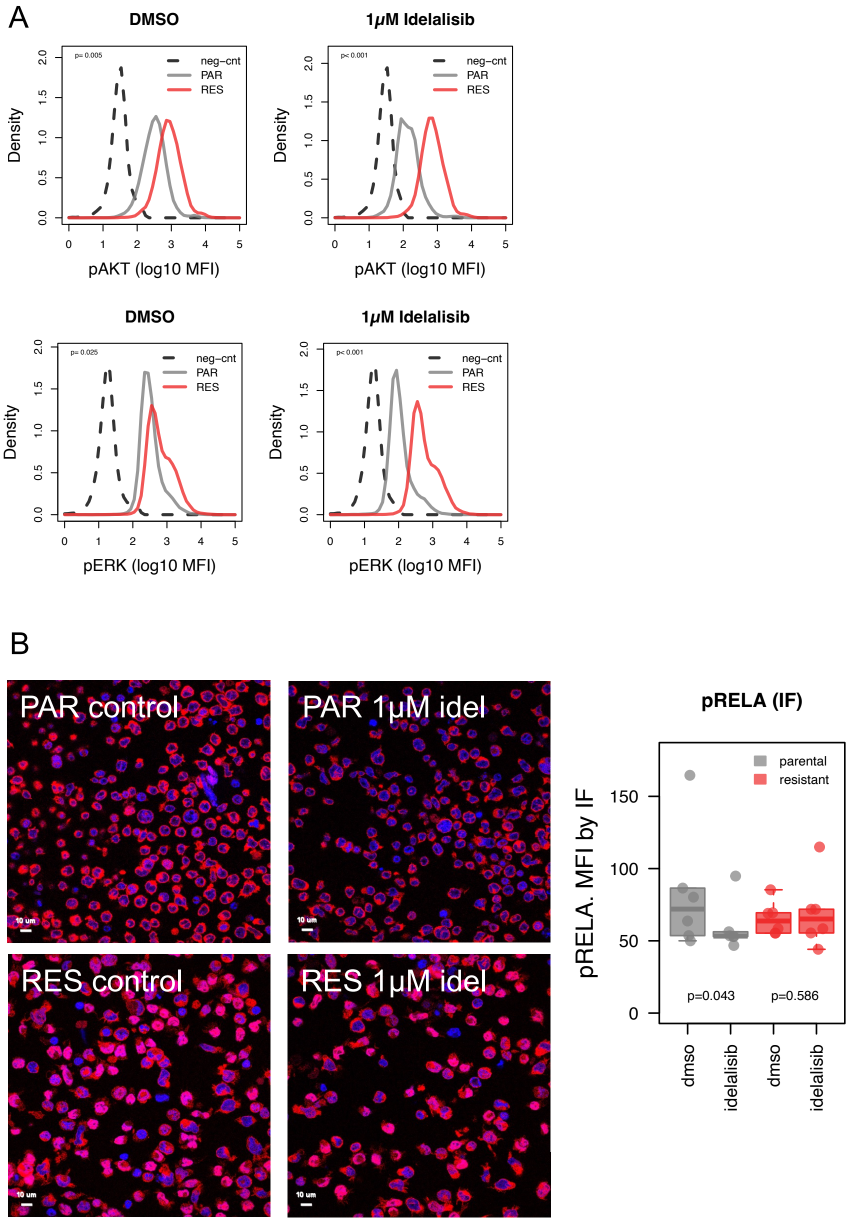


**Figure S8**. Levels of p-AKT (A, top) and p-ERK (B, bottom) were determined by phospho-flow cytometry upon 1μM of idelalisib. Density plots show the median MFI values of two replicates, negative control (dotted black), parental (grey), resistant (red). (B) Images on the left show the expression of pRELA by immunofluorescence upon 1μM of idelalisib (right) or control (DMSO, left) in parental and resistant (4hr exposure). Boxplot on the right represents the protein quantification from the immunofluorescence experiments. Data derived from two independent experiments. Three representative fields were evaluated from each independent experiment. P-values from Welch Two Sample test performed on mean MFI values.

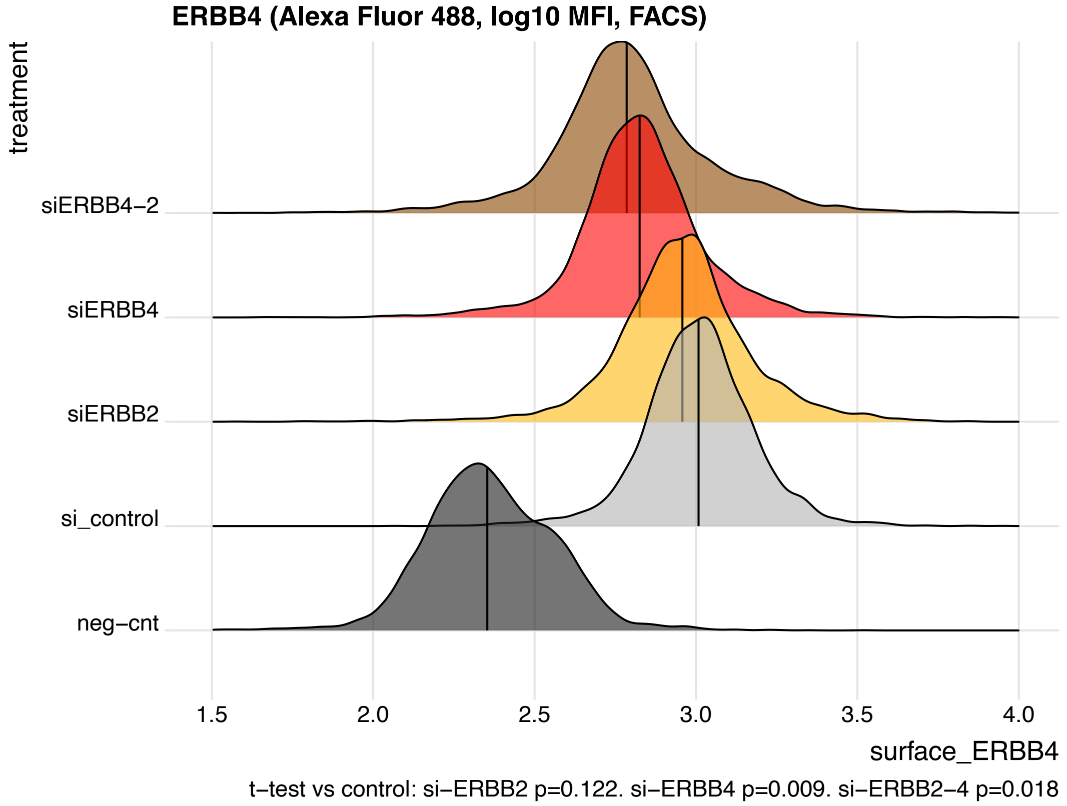


**Figure S9**. Ridgeplot on surface levels of ERBB4 by FACS in Karpas1718 resistant cells upon genetic silencing (siRNA) of ERBB2 (yellow), ERBB4 (red) or concomitant ERBB2 and ERBB4 (brown). Silencing control in grey and negative control in black. Data derived from two independent experiments. Statistically significant differences ($p<0.05$) were evaluated by t-test performed on mean MFI values.

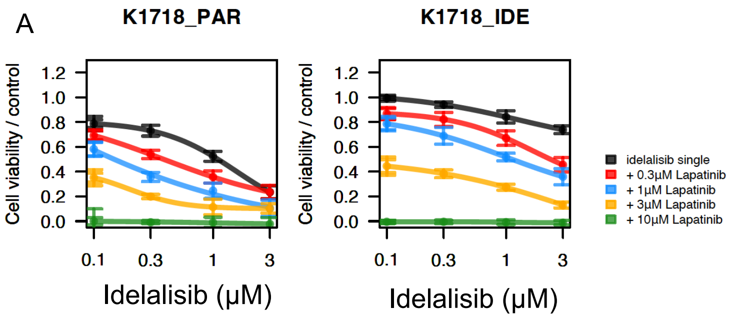


| p-value from moderated t-test comparing each combination to idelalisib single | | |
|---|---|---|
| Combination | Parental | Resistant |
| 0.3µM Lapatinib | 0.123 | 0.006 |
| 1µM Lapatinib | 0.014 | 0.002 |
| 3µM Lapatinib | 0.005 | <0.001 |
| 10µM Lapatinib | 0.002 | <0.001 |

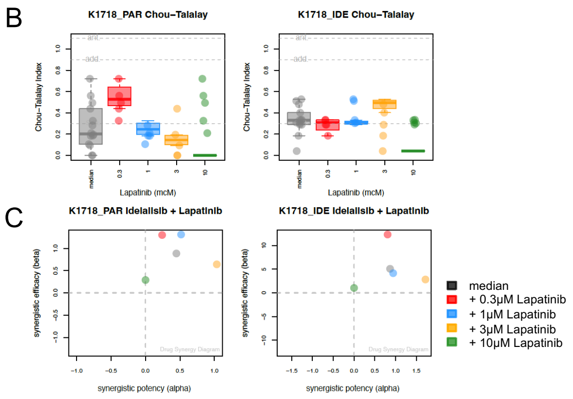


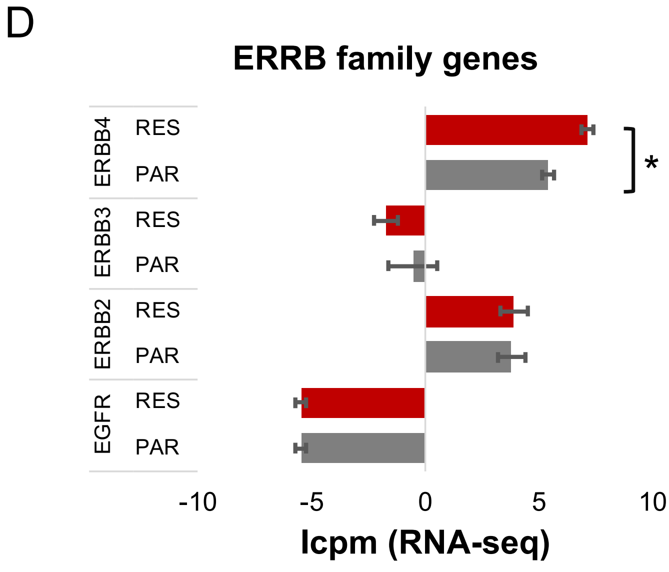


**Figure S10**. (A) Cell viability for the combination of idelalisib and the ERBB inhibitor lapatinib in parental and resistant by MTT assay (72h). Bars correspond to the mean of two independent experiments. Error bars represent standard deviation of the mean. Table contains p-values from a moderated t-test comparing each combination to idelalisib as single agent. The benefit of the combination was assessed both as synergism according to the Chou-Talalay combination index (B) (23) and as potency (x-axis) and efficacy (y-axis) according to the MuSyC algorithm (C) (3). K1718_PAR: parental, K1718_IDE: resistant. (D) Gene expression of ERBB family genes in parental (grey) and resistant (red) by RNA-seq. Bars correspond to the mean of two independent replicates. Error bars represent standard deviation of the mean. * for adjusted P-value lower than 0.05 from a moderated t-test.

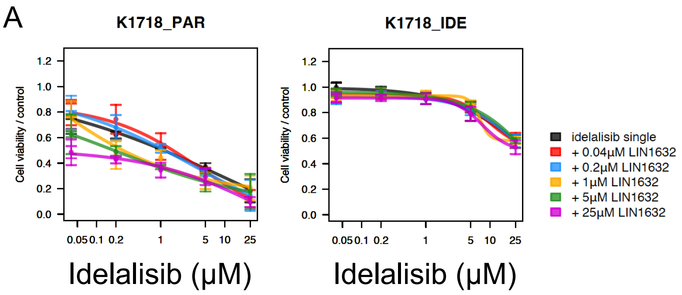


| p-value from moderated t-test comparing each combination to idelalisib single | | |
|---|---|---|
| Combination | Parental | Resistant |
| 0.04µM LIN1632 | 0.685 | 0.607 |
| 0.2µM LIN1632 | 0.965 | 0.319 |
| 1µM LIN1632 | 0.535 | 0.809 |
| 5µM LIN1632 | 0.224 | 0.829 |
| 25µM LIN1632 | 0.035 | 0.077 |

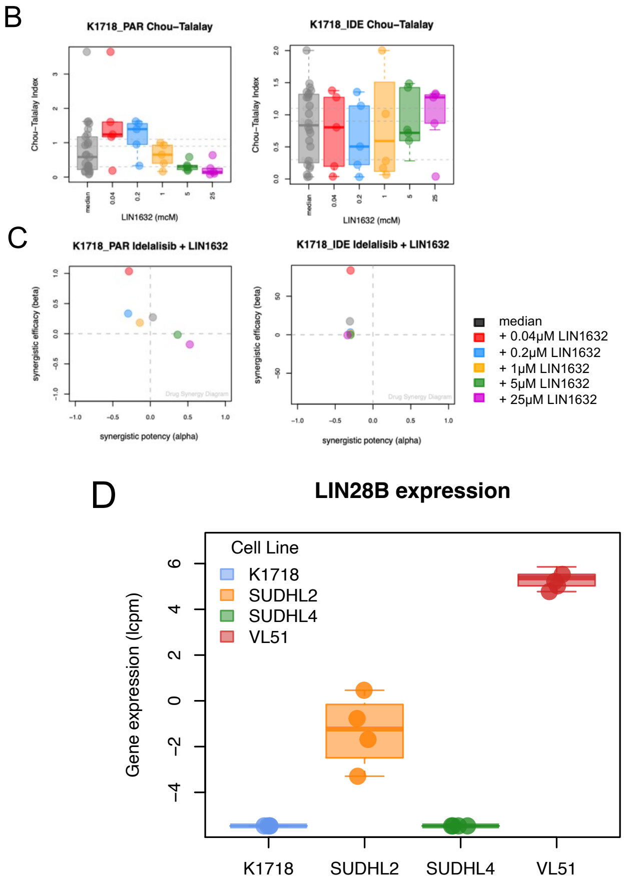


**Figure S11**. (A) Cell viability for the combination of idelalisib and the LIN28 inhibitor LIN1632 in parental and resistant by MTT assay (72h). Bars correspond to the mean of two independent experiments. Error bars represent standard deviation of the mean. Table contains p-values from a moderated t-test comparing each combination to idelalisib as single agent. The benefit of the combination was assessed both as synergism according to the Chou-Talalay combination index (B) (23) and as potency (x-axis) and efficacy (y-axis) according to the MuSyC algorithm (C) (3). K1718_PAR: parental, K1718_IDE: resistant. (D) Boxplot on the LIN28B expression (RNA-seq) in B-cell lymphoma cell lines: Karpas1718 (blue), SUDHL2 (yellow), SUDHL4 (green) and VL51 (red).

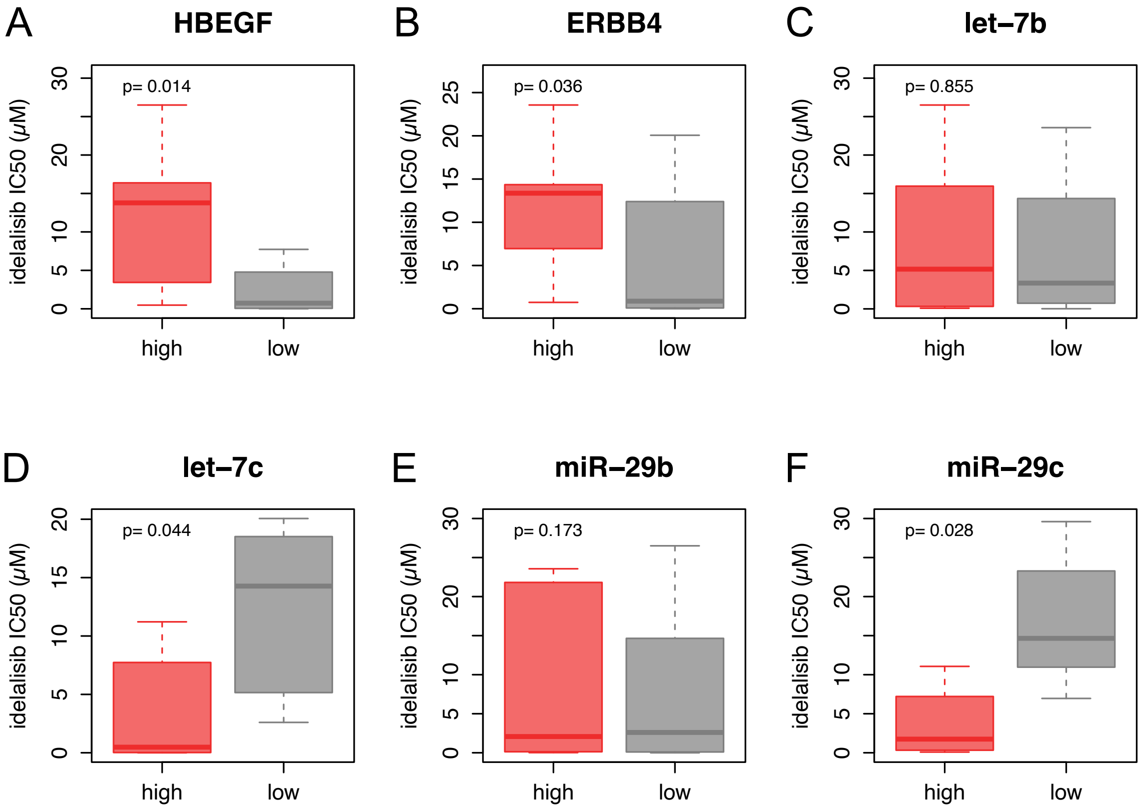


**Figure S12**. Expression levels of crucial factors are correlated with resistance to idelalisib. HBEGF (A) and ERBB4 (B) expression levels are inversely correlated with idelalisib sensitivity. Conversely expression of let-7c and miR-29c microRNAs is associated with sensitivity to idelalisib. Expression and sensitivity data were analyzed from a previous publication of our group in a panel of 34 B cell lymphoma cell lines (24). Cell lines were split in two groups based on higher or lower values than the median expression of the corresponding gene or miRNA. Mean of idelalisib IC50 were calculated for these two groups and compared by t-test.

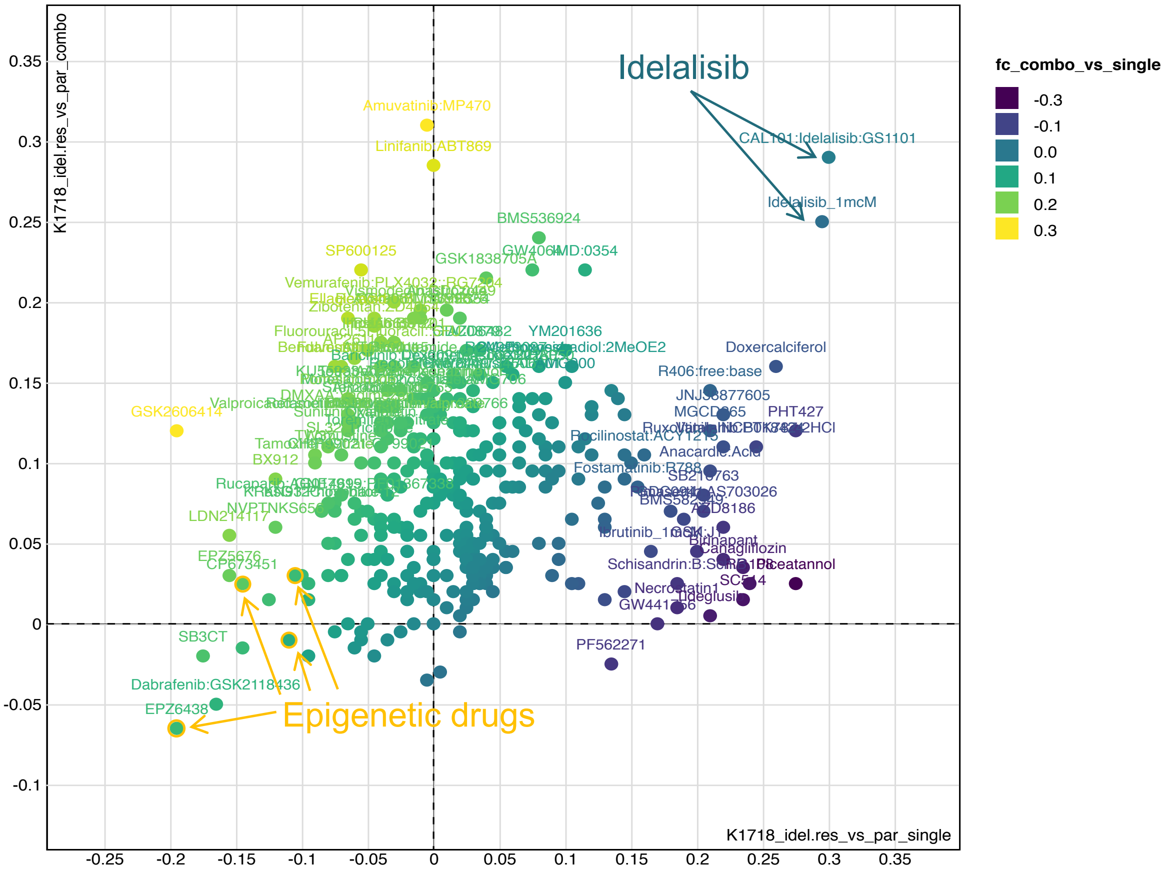


**Figure S13**. Resistant in parallel with parental clones of Karpas1718 cell line were exposed to a library of 348 compounds alone (single, 1µM) or in combination with 1µM idelalisib (combo) for 72h. Cell viability was measured by MTT assay (72h). Yellow for benefit in combo, purple benefit in single treatment.

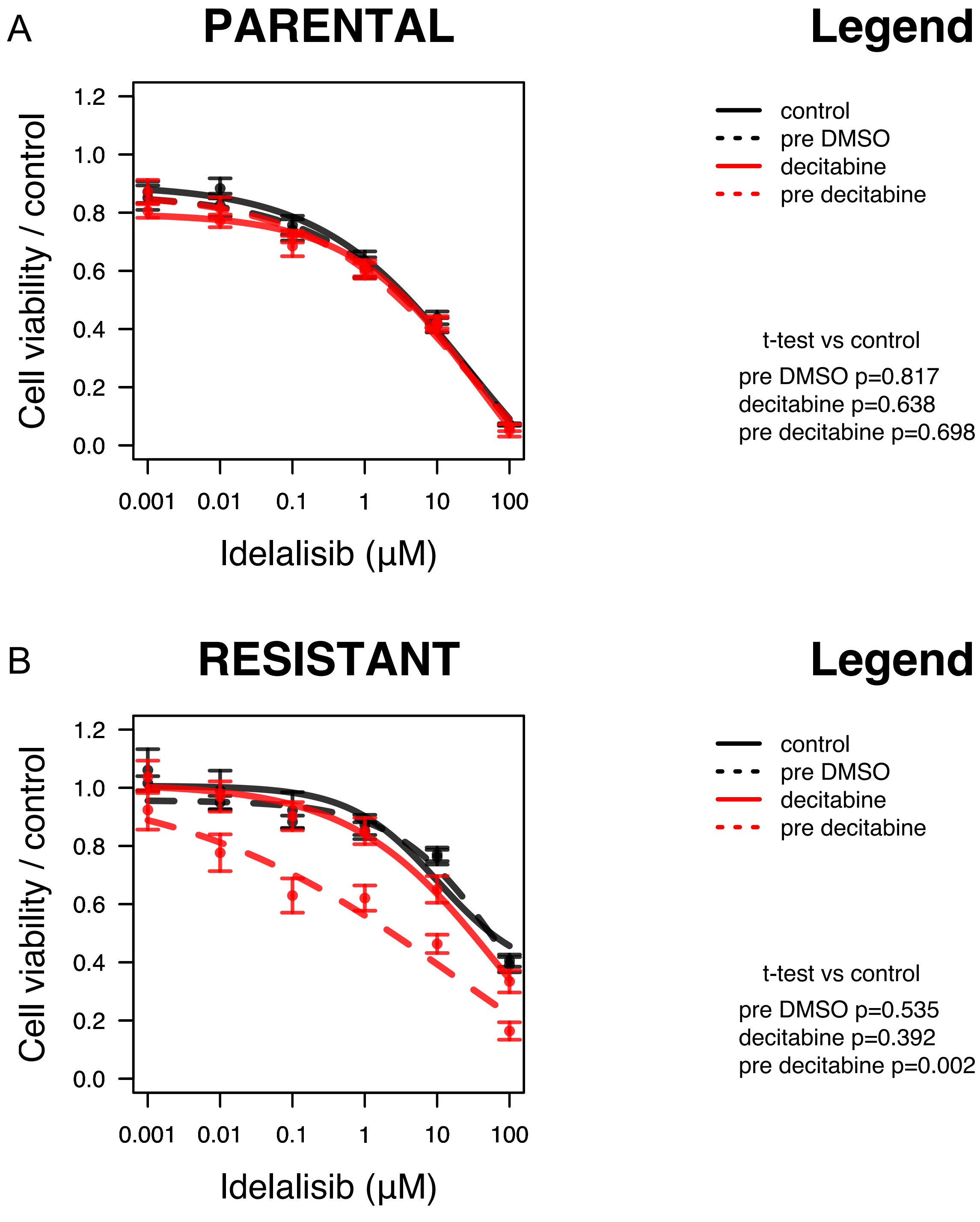


**Figure S14**. Cell viability was evaluated by MTT assay (72h). Karpas1718 parental (A) or resistant (B) were exposed to DMSO (control, black) or decitabine (100nM, red), given concomitantly (continuous lines) or 72 hours before idelalisib (pre DMSO in dashed black or pre decitabine in dashed red). Error bars correspond to standard deviation of the mean. Data derived from two independent experiments. Statistically significant differences were evaluated by moderated t-test comparing each treatment to control (DMSO).

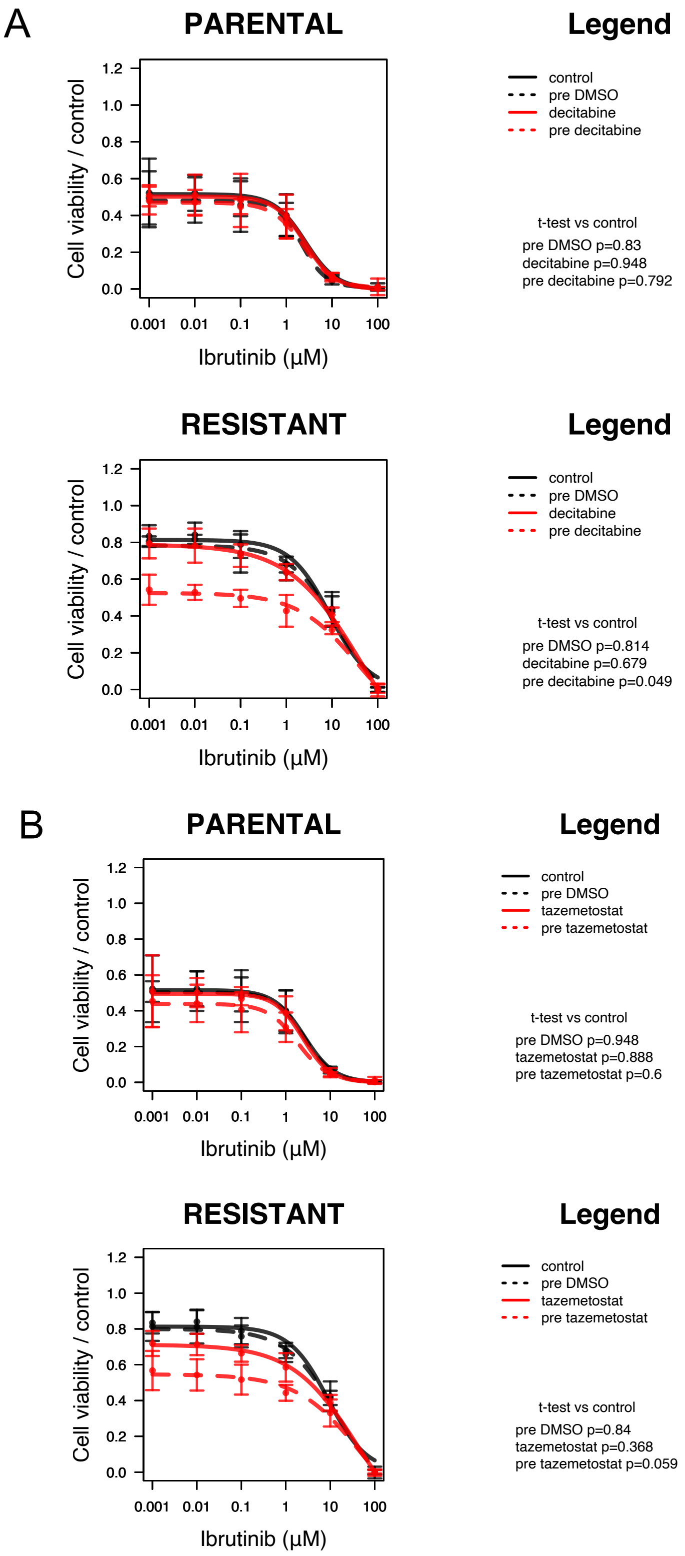


**Figure S15**. Sensitivity to ibrutinib was evaluated by MTT assay (72h). Karpas1718 parental (top) or resistant (bottom) were exposed to 100nM decitabine (A), 5µM tazemetostat (B), or DMSO (control, black) given concomitantly (continuous lines) or 72 hours before ibrutinib (pre DMSO in dashed black or pre decitabine/tazemetostat in dashed red). Error bars correspond to standard deviation of the mean. Data derived from two independent experiments. Statistically significant differences were evaluated by moderated t-test comparing each treatment to control (DMSO).

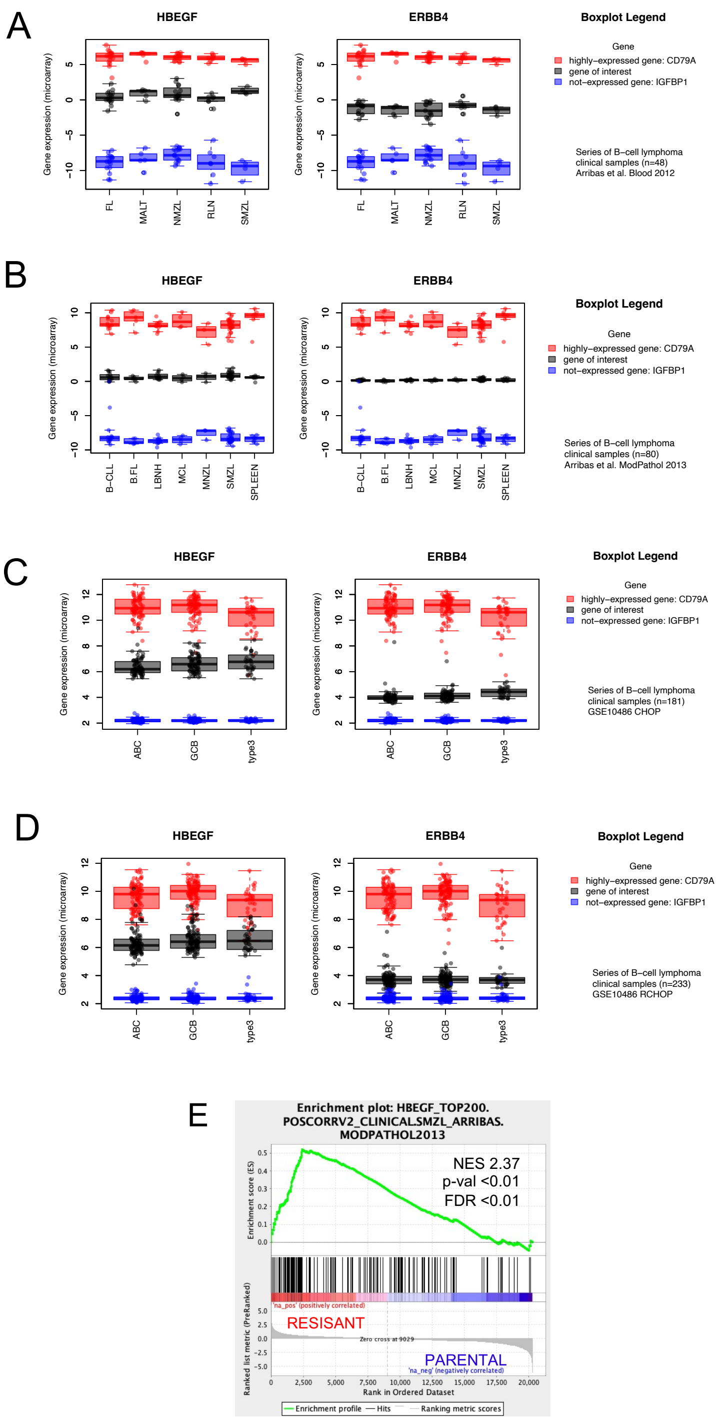


**Figure S16**. Factors associated with resistance to idelalisib in cell lines are expressed in clinical specimens. Expression levels of genes related to idelalisib resistance were studied across different subtypes of B cell lymphoma: (A) n=48 (25), (B) n=80 (26). B-CLL: chronic lymphocytic leukemia, B.FL: follicular lymphoma, LBNH: non-specified non-Hodgkin B cell lymphoma, MALT: MZL of the mucosa associated tissue, MCL: Mantle cell lymphoma, NMZL: nodal marginal zone lymphoma, SMZL: splenic marginal zone lymphoma. Expression levels of genes related to idelalisib resistance were studied across the subtypes of diffuse large B cell lymphoma (DLBCL, two series: (C) n=181 and (D) n=223 from GSE10846 (27). ABC: activated B cell like DLBCL, GCB: germinal center B cell like DLBCL, type3: type3 DLBCL. Red for highly-expressed gene in B cells (CD79A), blue for not-expressed gene in B cells (IGFBP1), black for the gene of interest. (E) Gene set enrichment analyses comparing resistant versus parental for the top-200 genes positively correlated genes with *HBEGF* in SMZL clinical specimens (26). NES: normalized enrichment score, p-val: nominal p-value, FDR: false discovery rate.
